# The Nematicide Tioxazafen Disrupts Proteasome Function via Cytochrome P450 Bioactivation

**DOI:** 10.64898/2026.07.10.737828

**Authors:** Lucia Chen, Kaiden Thompson, Muntasir Kamal, Kateryna Sihuta, Irini Topalidou, Andrew R. Burns, Nazli H. Farshour, Brittany Cooke, Jessica Knox, Yanqing Jiang, Modhar Al Qasser, Ermira Shuteriqi, Matej Usaj, Justin Ching, Arthur Flaget, Michael Costanzo, Guihong Tan, Jessica Lacoste, Mark Lautens, Brenda Andrews, Charles Boone, Mikko Taipale, Nicolas J. Lehrbach, Peter J. Roy

## Abstract

Tioxazafen is an effective nematicide whose commercialization was halted because handlers reportedly developed rashes after working with seeds coated with a tioxazafen-laced cocktail. Here, we show that tioxazafen is bioactivated into toxic products by nematode and human cytochrome P450s. Through systematic analyses, we found that bioactivated tioxazafen disrupts proteasome function, leading to the accumulation of the NRF1 transcription factor ortholog SKN-1A in the nematode *C. elegans*, and a ‘bounce-back’ transcriptional up-regulation of proteasome components. Genetic upregulation of the *C. elegans* proteasome supresses tioxazafen’s lethality, indicating that proteotoxicity is a key contributor to death. A survey of human P450s revealed that skin-expressed CYP1A1 toxifies tioxazafen and may account for the reaction to tioxazafen-coated seeds. We also found that rabbit CYP1A1 fails to bioactivate tioxazafen, which may explain the pre-market failure to detect robust adverse skin reactions. Our work highlights vulnerabilities in pre-market toxicological assays and a provides potential solution to prospectively identifying P450 toxication events.

**One-Sentence Summary:** Tioxazafen is bioactivated by cytochrome P450s into a proteasome disruptor.

## Introduction

Plant parasitic nematodes (PPNs) are particularly pervasive and pestiferous beasts. They are estimated to destroy upwards of $200 billion USD in crops globally every year (Nicol *et al*. 2011; Dyrdahl-Young *et al*. 2020; Khanal and Land 2023) and farmers collectively spend over $1 billion USD annually to control them (MARKETS 2024; Research 2024). The primary means of PPN control are small molecule nematicides because they are relatively inexpensive and have historically been effective. Over the past couple of decades however, multiple classes of nematicides have been banned or severely restricted because of off-target effects, thereby limiting the tools in our arsenal to control PPN infestations (Costa *et al*. 2008; Donley 2019; Desaeger *et al*. 2020). The dearth of effective nematicides has inspired many groups, including ours, to develop new selective nematicides (Lahm *et al*. 2017; Harrington *et al*. 2022; Burns *et al*. 2023; Davie *et al*. 2024; Knox *et al*. 2024; Elfawal *et al*. 2025; Fahs *et al*. 2025).

In the mid 2000s a novel nematicide called tioxazafen (3-phenyl-5-(2-thienyl)-1,2,4-oxadiazole) was developed and subsequently acquired by Monsanto Inc (Slomczynska *et al*. 2015; Faske *et al*. 2022). Tioxazafen showed promise because it has nematode-selective activity against a wide-range of devastating plant parasitic nematodes (Slomczynska *et al*. 2015; South *et al*. 2019; Faske *et al*. 2022). Tioxazafen was formulated as part of a multi-compound seed treatment for corn, cotton and soybean, and was commercialized as NemaStrike. Unfortunately, Nemastrike was pulled from commercialization efforts in 2017 and again in 2019, following Bayer AG’s acquisition of Monsanto, reportedly due to skin irritation in handlers (Polansek 2017; Smith 2019). The mechanism of action of tioxazafen was not well-understood (South *et al*. 2019) and data linking the tioxazafen molecule itself to the skin irritation are lacking.

Previously, our group identified the pro-nematicides selectivin (Burns *et al*. 2023) and cyprocide (Knox *et al*. 2024). Both selectivin and cyprocide are metabolized by distinct cytochrome P450 enzymes in the nematode worm *C. elegans* into reactive electrophilic products that bind low molecular weight thiols, including glutathione, γ-glutamylcysteine, and cysteine. We demonstrated that CYP-35C1 and CYP-35D1 were necessary for selectivin and cyprocide bioactivation in *C. elegans*, respectively. Through heterologous expression in the yeast *Saccharomyces cerevisiae*, we also demonstrated that the aforementioned *C. elegans* P450s were sufficient for the bioactivation of the respective pro-nematicides. Using this yeast assay, we also identified a cytochrome P450 from the plant parasitic nematode *Meloidogyne incognita* that can carry out the bioactivation reactions (Burns *et al*. 2023; Knox *et al*. 2024). We speculate that the bioactivated molecules’ lethality stems from the reactive metabolites i) consuming redox buffers like glutathione and in turn increasing redox stress, and ii) binding essential proteins via cysteine interactions and disrupting their function. This interpretation of the mechanism of killing by reactive electrophilic metabolites is widely accepted (Guengerich 2021).

Here, we show that tioxazafen is also a pro-nematicide but has a distinct mechanism of bioactivation compared to selectivin and cyprocide. Systematic genetic interaction analyses, systematic stress reporter screens, RNAseq and other assays reveal that bioactivated tioxazafen ultimately inhibits *C. elegans* proteasome function while coincidentally repressing the transcription of several molecular machines involved in cell cycle progression. The proteasome is a highly conserved multi-subunit complex that is a key component in regulating proteostasis within eukaryotic cells (Bard *et al*. 2018) and is key in regulating progression through the cell cycle (Jang *et al*. 2020).

We used our previously-established yeast-based P450 expression system (Burns *et al*. 2023; Knox *et al*. 2024) to survey the three families (1-3) of P450s commonly associated with xenobiotic metabolism in humans (Rendic and Guengerich 2015). We found that human CYP1A1, and to a lesser extent CYP1B1, are sufficient to toxify tioxazafen in yeast and that CYP1A1 robustly does so in HEK293T cells. RNAseq of CYP1A1-expressing HEK293T cells incubated in tioxazafen revealed markers of proteotoxic stress. Both CYP1A1 and CYP1B1 are expressed in skin (Katiyar *et al*. 2000; Chen *et al*. 2024), providing a reasonable explanation for why tioxazafen-laced products reportedly induce rashes in humans (Polansek 2017; Smith 2019). Finally, we show that rabbit CYP1A1 and rat CYP1B1 cannot toxify tioxazafen despite being functional in our assay. Our data suggest that pre-market animal safety tests can mislead the assessment of product safety because of sequence divergence of xenobiotic defense mechanisms.

## Results

### Tioxazafen Killing of *C. elegans* is Cytochrome P450-Dependent

Previously, we screened our collection of candidate nematicides (Burns *et al*. 2015) for those whose lethality might be dependent on cytochrome P450s (P450s) (Knox *et al*. 2024). To do this, we disrupted the *C. elegans* EMB-8 P450-oxidoreductase (aka, POR or CPR) co-factor that is necessary for microsomal P450 activity (Rappleye *et al*. 2003; Coon 2005; Harlow *et al*. 2018). We reasoned that nematicides that lose potency upon disruption of *emb-8* are likely to be P450-bioactivated. In our screen, we identified two structural analogs of tioxazafen called wact-2, which is brominated, and wact-180, which is trifluoromethylated (Knox *et al*. 2024). A dose-response analysis of tioxazafen and analogs showed that their lethality is indeed *emb-8*-dependent (Fig. 1a). We previously used tioxazafen as a control in our assays and have demonstrated that it effectively kills *C. elegans* larvae and dauers, as well as the embryos of the plant parasitic nematode *Meloidogyne hapla* (see Fig. 1e of (Knox *et al*. 2024)).

**Figure 1.**
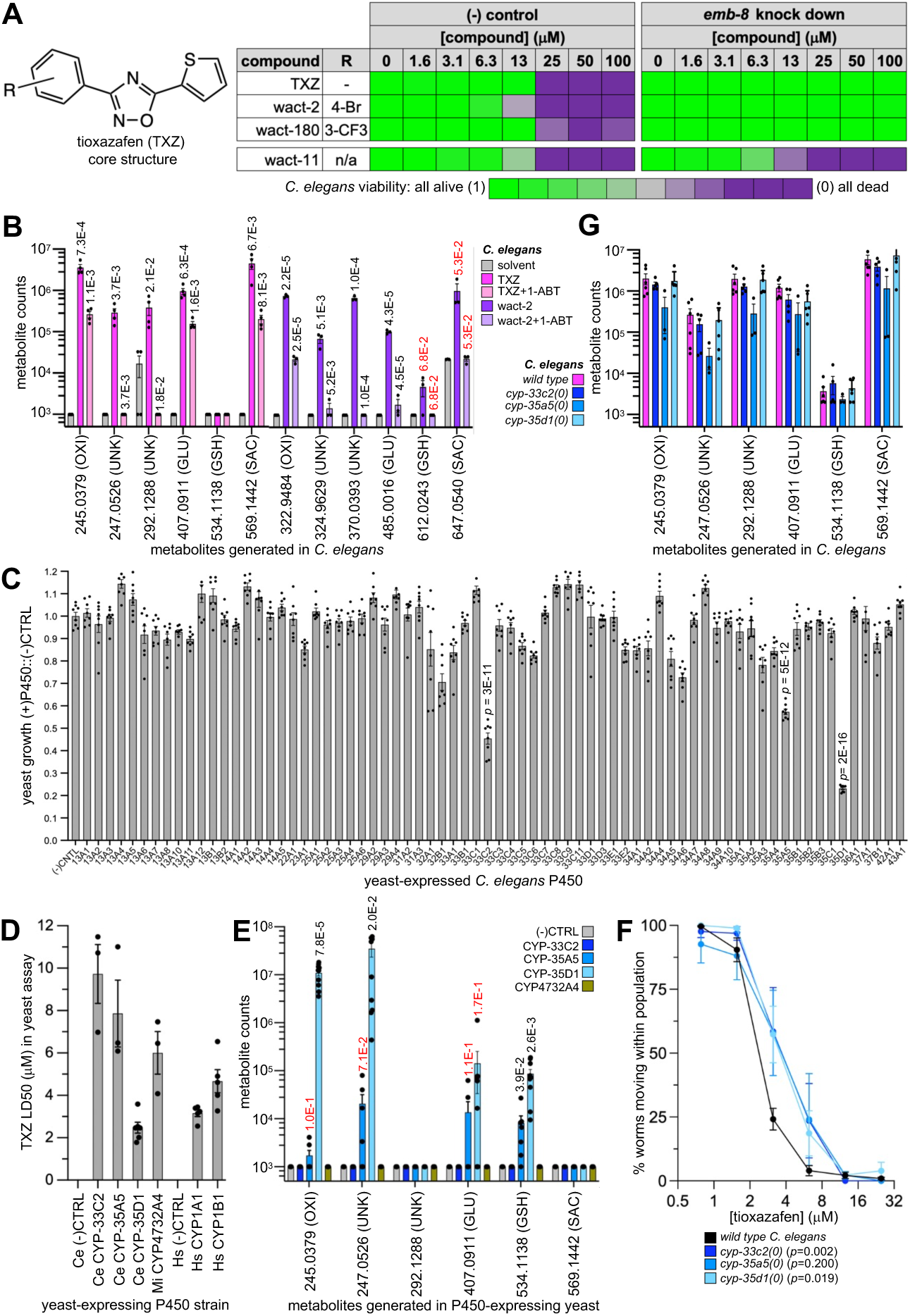
Tioxazafen is Toxified by Cytochrome P450s. **A.** Tioxazafen and analog lethality is dependent on the *C. elegans* cytochrome P450 oxidoreductase EMB-8. Note that the viability scale is relative to animals grown in DMSO solvent control-only. The negative control is the succinate dehydrogenase inhibitor wact-11, which is not EMB-8 dependent. **B.** Tioxazafen and wact-2-derived metabolites generated in *C. elegans*. The exact mass of TXZ is 229.0430 and wact-2 is 306.9535. The Students’ T-test *p* values shown atop each bar is in comparison to the data in the previous bar. **C.** A PIXY survey of the ability of the indicated yeast-expressed *C. elegans* P450 to toxify 100 μM tioxazafen. n=8; all *p* values less than 1E-10 relative to the negative control using a two-tailed Student’s T-test are shown. **D.** Tioxazafen LD50 analyses for yeast expressing the indicated P450. Ce, *C. elegans*, Mi, *Meloidogyne incognita*, Hs, *Homo sapiens*. N≥3, n=2. Statistical comparisons relative to the (-) control was not possible given that LD50s could not be established for the (-) within the concentration range tested (i.e., the LD50 of the (-) control with TXZ is >>100 μM). **E.** Tioxazafen-derived metabolites generated in yeast expressing the indicated P450. N≥3, n=2. The Students’ T-test *p* values shown atop each bar is in comparison to the (-) control. **F.** Three-day viability assay of the indicated likely loss-of-function mutant relative to wild type. The x-axis is Log_2_. N≥5 independent biological trials, each with n≥50 worms per trial per concentration. The *p* value of the EC50s, as determined by a Mann-Whitney test is indicated in the legend. **G.** The generation of TXZ-derived metabolites in the three *C. elegans* P450 mutants relative to the wild type control. N≥3, n=2. A Students’ T-test was performed relative to the wild type control and no significant differences were observed (*p*>0.05). *p* values In B and E, *p* values are coloured red if *p*<0.05. In B, E and G, exact masses are shown on the x-axis along with the likely type of conjugate in brackets (OXI, oxidized; UNK, unknown; GLU, glucosidated; GSH, glutathione; SAC, disaccharide). In all graphs, the standard error of the mean is shown.

If tioxazafen is P450-bioactivated, we would expect to see metabolic derivatives of tioxazafen in worm lysates using liquid chromatography-coupled mass spectrometry (LCMS). We observed multiple metabolites that were present in drug-treated worms but not in solvent only-controls (Fig. 1b; Supplemental Figs.1&2). We did the same analysis with wact-2 and observed similar metabolites as those seen with tioxazafen (accounting for the difference in the parent masses) (Fig. 1b). Because wact-2 is brominated (and bromine has two abundant naturally occurring isotopes that make the recognition of brominated metabolites definitive by LCMS analyses) we could be confident that these metabolites are derivatives of the parent structures. Finally, we also tested whether the broad-spectrum P450 inhibitor 1-ABT could suppress metabolite formation and found that it could (Fig. 1b). Together, these results are consistent with the idea that tioxazafen is metabolized by P450s into a lethal metabolite.

Tioxazafen and its bioactive analogs harbour a thiophene ring, which can be metabolized by P450s into reactive sulfoxidated or epoxidated structures (Gramec *et al*. 2014). Indeed, the oxidized mass (245.0379 [H+]) of tioxazafen is consistent with these and similar reactions (Podgorski *et al*. 2023) (Fig. 1b; Supplementary Figs. 1&2). However, unlike the metabolism of selectivin and cyprocide (Burns *et al*. 2023; Knox *et al*. 2024), we found little evidence of abundant tioxazafen conjugates with low molecular weight (LMW) thiols such as glutathione in *C. elegans*. Consistent with this observation, we were not able to rescue tioxazafen or wact-2-induced lethality with NACET, which is converted into excess LMW thiols in the cell (Giustarini *et al*. 2012), and is able to rescue the lethality induced by bioactivated selectivin and cyprocide (Knox *et al*. 2024) (Supplemental Fig. 3). We conclude that the nature of tioxazafen’s bioactivated product(s) is generally distinct from that of selectivin or cyprocide.

### *C. elegans* CYP-35D1 Is Sufficient for Tioxazafen’s Lethality

To identify the cytochrome P450s responsible for tioxazafen bioactivation, we systematically tested all 73 *C. elegans* microsomal P450s for their ability to toxify tioxazafen when heterologously expressed in the yeast *S. cerevisiae*. We call this assay the P450 Interaction eXperiment in Yeast (PIXY). We expressed the *C. elegans* POR (EMB-8) from a *GAL1* promoter integrated into the *CAN1* locus and the P450s or the empty vector control from the *GAL1* promoter from a low-copy plasmid (see methods). From the initial PIXY survey, we identified three P450s that robustly toxify tioxazafen; CYP-33C2, CYP-35A5 and CYP-35D1 (*p*<1E-10) (Fig 1c). Dose-response analysis confirmed the interactions and showed that CYP-35D1 most potently bioactivates the nematicide (Fig 1d). A similar PIXY survey of 19 P450s from the plant parasitic nematode *Meloidogyne incognita* revealed CYP4732A4 to toxify tioxazafen (Fig 1d; Supplemental Fig. 4). We used LCMS to show that the yeast strains were capable of generating some of the same exact masses of tioxazafen metabolites as observed in worms (Fig 1e). *C. elegans* knockouts of each of the three P450s failed to exhibit robust resistance to the molecule in larval viability assays with only minor shifts in the EC50s (*p*<0.05 for two of the three mutants) (Fig 1f). The lack of robust resistance is likely due to redundancy among the P450s in being able to generate toxic metabolites, which is consistent with our inability to isolate mutants that resist tioxazafen’s lethality from a screen of over a quarter of a million mutant genomes (Burns *et al*. 2015). We also failed to see significant differences in the production of metabolites in the mutant backgrounds relative to the wild type controls (Fig 1g), which is consistent with the phenotypic data. We conclude that multiple nematode P450s can toxify tioxazafen.

### Bioactivated Tioxazafen Genetically Interacts with Mutant Proteasome Components in Yeast

We took several approaches to understand the impact of bioactivated tioxazafen on living systems. First, we systematically identified genetic enhancers of bioactivated tioxazafen in yeast using synthetic genetic array (SGA) technology (Costanzo *et al*. 2016). A yeast strain harboring SGA reporters and expressing either the plasmid expressing CYP-35D1 or an empty-vector negative control plasmid were crossed into 5193 strains harbouring either gene deletions or temperature-sensitive (ts) alleles of 4679 unique nonessential and essential yeast genes. We then grew these strains on a concentration of tioxazafen (15 μM) that slows the growth of CYP-35D1-expressing cells by 30% on solid substrate and then asked which mutant genes enhance the growth defect of the CYP-35D1-expressing cells relative to the growth rate of cells harbouring only the (-) control. The logic of the experiment is that processes that are antagonized by bioactivated tioxazafen should show heightened sensitivity to low concentrations of the bioactivated compound when the processes are already compromised by mutation (Piotrowski *et al*. 2017). 61 genes were found to enhance the lethality of bioactivated tioxazafen (*p*<1E-02 in each of three independent trials) (Fig. 2a; Datafile 1).

**Figure 2.**
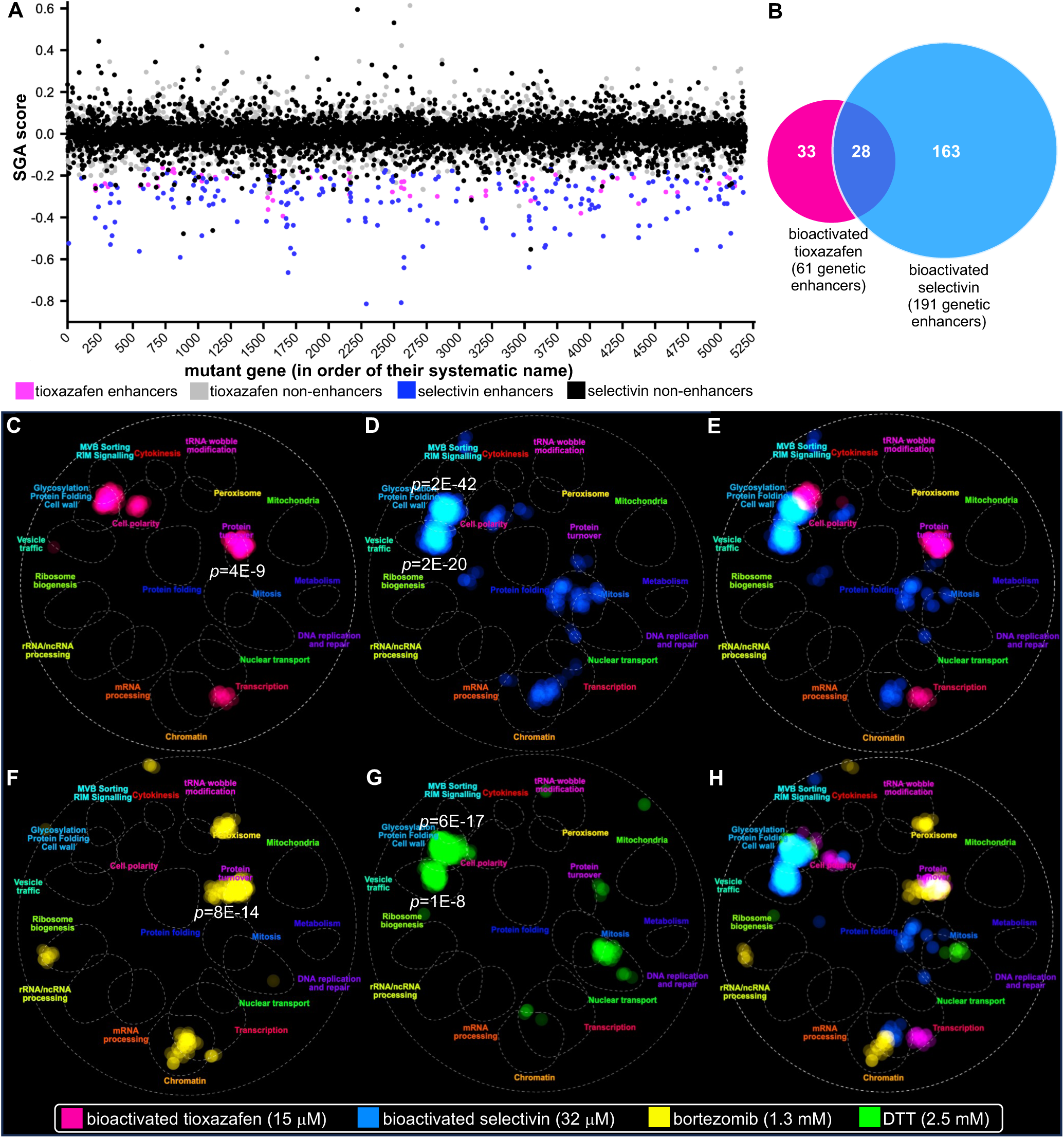
Bioactivated Tioxazafen Disrupts Protein Turnover in Yeast. **A.** A summary of the SGA screens for genetic modifiers of bioactivated tioxazafen and bioactivated selectivin. Shown is the average SGA score of each genetic disruption relative to the EC30 concentration of tioxazafen and selectivin over three independent trials with each trial having an n of 4 technical repeats. A low SGA score indicates that the mutant gene enhanced the growth defect of the strain expressing the relevant nematode P450 relative to the empty-vector control mutant strain. A high SGA score shows the opposite (i.e., genetic suppressors of the bioactivated compound). **B.** A Venn diagram showing the overlap of the yeast genetic enhancers, identified through SGA, of bioactivated tioxazafen and bioactivated selectivin (*p*<0.01 in each of three independent trials with each trial having an n of 4). **C-H**. CellMap enrichment plots of the genetic enhancers of bioactivated tioxazafen (C), bioactivated selectivin (D), the overlap between tioxazafen and selectivin (E), bortezomib (F), DTT (G), and the overlap between tioxazafen, selectivin, bortezomib and DTT. The enrichment value for all processes enriched in hits with *p*<E-07 is shown.

To understand how selective these enhancers are for bioactivated tioxazafen, we performed a second SGA analysis focused on the selectivin nematicide that is bioactivated by the nematode P450 CYP-35C1 (Burns *et al*. 2023). We introduced a plasmid expressing CYP-35C1 into the yeast mutant strain collections described above and used a concentration of selectivin (32 μM) that slows the growth of CYP-35C1-expressing yeast cells by 30% on solid substrate. We identified 191 genes that enhance bioactivated selectivin lethality (*p*<0.01 in each of three independent trials) (Fig. 2a; Datafile 1). A Venn diagram illustrates that 33 of the 61 genetic enhancers are unique to bioactivated tioxazafen (Fig. 2b).

To identify and display which processes are enriched for these enhancers, we performed a Spatial Analysis of Functional Enrichment (SAFE) (Baryshnikova 2016) using the online Cell Map tool (Usaj *et al*. 2017). Previous genome-wide analyses generated a complete genetic interaction network revealing the functional organization of a yeast (Costanzo *et al*. 2016). SAFE identifies regions of the global yeast genetic network corresponding to unique cellular functions that are enriched for a particular set of genes, such as the enhancers identified in our PIXY screens. The process most significantly enriched with enhancers that selectively bioactivate tioxazafen was protein turnover (*p*<4E-9) (Fig. 2c); no other process was enriched with enhancers below *p*<1E-5 (Supplemental Table 1). By contrast, the mutant genes that enhance the lethality of bioactivated selectivin are enriched in glycosylation and protein folding (*p*<2E-42) followed by those involved in vesicle trafficking (*p*<2E-20) (Fig. 2d; Supplemental Table 1) and are largely distinct from the processes affected by bioactivated tioxazafen (Fig. 2e). Enrichment of similar groups is seen with traditional GO analyses (DataFile 1).

Given that the proteasome is a key regulator of protein turnover, we asked whether genetic interactors of the proteasome inhibitor bortezomib (Costanzo *et al*. 2021) overlap those of bioactivated tioxazafen, and found that they do (Fig. 2f and 2h). This observation raised the possibility that bioactivated tioxazafen may disrupt the proteasome in yeast. By contrast, a hypothesis-driven search revealed that the pattern of genetic interactions with bioactivated selectivin is remarkably similar to that of the reducing agent dithiothreitol (DTT) (Berry *et al*. 2011) (Fig. 2g-2h). We reason that the ability of bioactivated selectivin to react with low molecular weight thiols (Burns *et al*. 2023) allows the bioactivated molecule to also react with the sulfur groups on cysteine-containing proteins and disrupt disulfide-bond formation, which is a key outcome of DTT treatment (Konigsberg 1972; Braakman *et al*. 1992).

### Bioactivated Tioxazafen Induces Proteasome Stress Likely Through Proteasome Inhibition

To understand how our candidate nematicides (Burns *et al*. 2015) affect *C. elegans*, we systematically investigated the impact of our 478 worm-active (aka ‘wactives’) small molecules on the expression of 12 different transcriptional GFP stress reporters at the end of a 24-hour incubation. These strains harboured reporters of oxidative stress (*gst-4p*::GFP and *sod-3p*::GFP (Libina *et al*. 2003; Choe *et al*. 2009)), reporters of infection (*cnc-2p*::GFP, *irg-1p*::GFP, and *nlp-29p*::GFP (Pujol *et al*. 2008; Dunbar *et al*. 2012)), a reporter of osmotic stress (*gpdh-1p*::GFP (Lamitina *et al*. 2006)), several reporters of the unfolded protein response (*hsp-4p*::GFP, *hsp-6p*::GFP, *hsp-16.41p*::GFP; *hsp-70p*::GFP, and *hsp-90p*::GFP (aka *daf-21p*::GFP) (Yoneda *et al*. 2004; Hartwig *et al*. 2009; Van Oosten-Hawle *et al*. 2013; Liu *et al*. 2014)), and a reporter of the proteasome (*rpt-3p*::GFP) that is up-regulated in response to proteasome dysfunction (Lehrbach and Ruvkun 2016). We displayed the resulting data by ordering the molecules according to structural similarity along the y-axis. A distinct and prominent pattern emerged from the analysis whereby the cluster of 15 tioxazafen-family members elicited strong expression from the osmotic, unfolded protein response (UPR), and proteasome reporters (Fig. 3a). The UPR and proteasome reporter results with tioxazafen were confirmed using RNAseq (see below; Supplemental Fig. 5).

**Figure 3.**
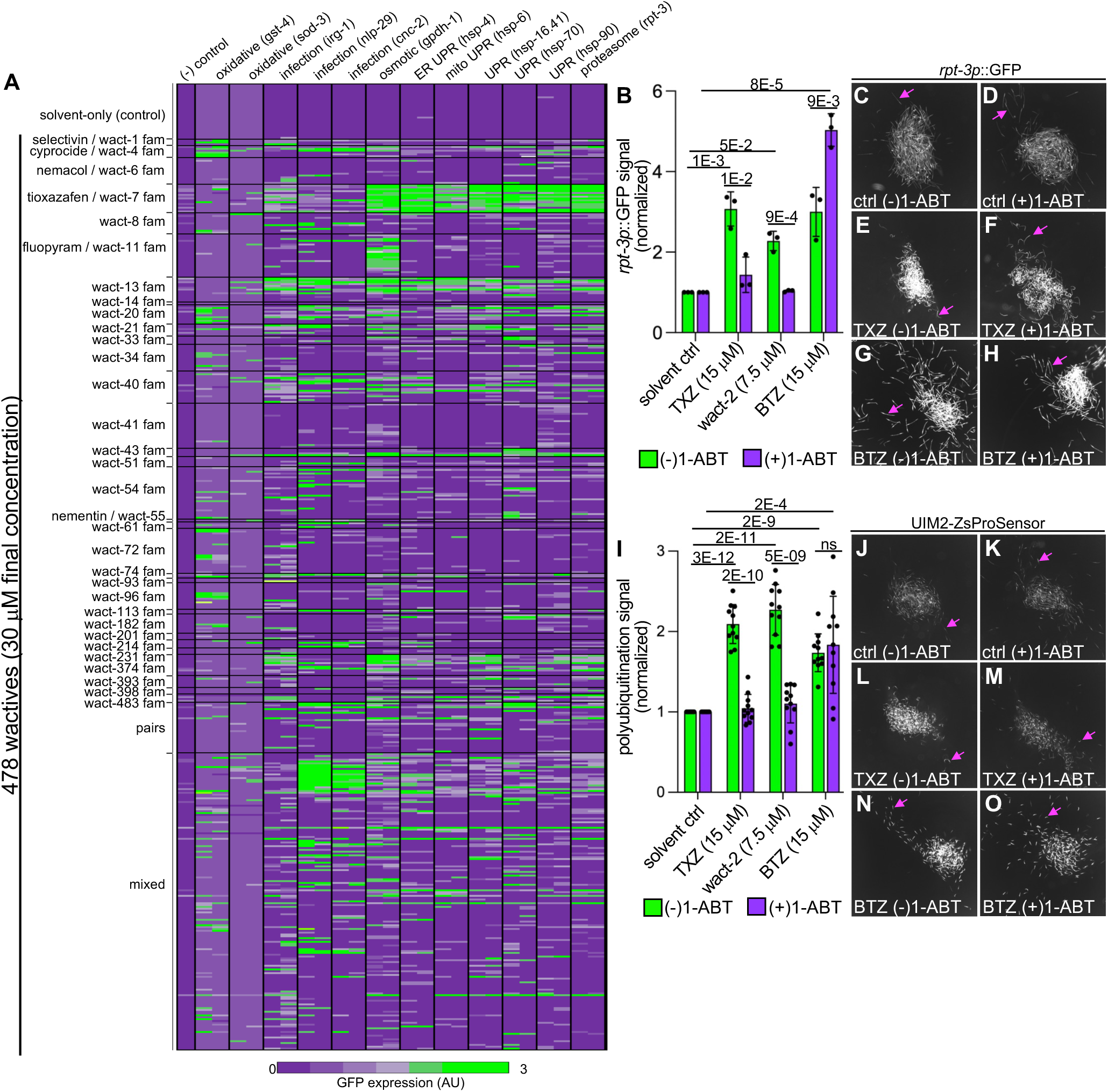
Tioxazafen and Structural Analogs Up-Regulate the Expression of a Proteasome Stress Reporter and Others. **A.** A systematic screen of our wactive library, where each compound is given a ‘wact’ designation. Molecules along the y-axis are ordered according to structural families (fam). Each assay was performed in duplicate with ∼20 animals per well (n=2). **B.** A detailed analysis of the expression of the *rpt-3p*::GFP proteasome stress reporter in response to sub-lethal concentrations of tioxazafen (TXZ), wact-2, or bortezomib with and without the cytochrome P450 inhibitor 1-ABT. Three independent trials (N=3) with at least 6 technical repeats per trial (n=6), with ∼150 worms per well in each technical repeat, was performed. **C-H.** Examples of the *rpt-3p*::GFP fluorescence observed in the wells from selected experiments from ‘B’. The pink arrow highlights a single worm in each of the wells. **I.** An analysis of the accumulation of polyubiquitinated proteins using the YD90 vha-6p::UIM2-ZsProSensor strain. The experiment was done in same manner as that described for B. **J-O.** Examples of the UIM2-ZsProSensor fluorescence accumulated in the YD90 strain in the indicated conditions of selected experiments from ‘I’. The pink arrow highlights a single worm in each of the wells. In each graph, the standard of deviation is shown, as are the relevant p values calculated from a two-sided Student’s T-test.

Through additional experimentation, we confirmed that the transcriptional *rpt-3* proteasome reporter is upregulated in response to both tioxazafen and wact-2 (Fig 3b-3h). This upregulation can be suppressed by the pan-P450 inhibitor, 1-ABT (Fig 3b and 3f), indicating that P450-bioactivated tioxazafen induces proteasomal stress. As positive control, we analyzed *rpt-3p*::GFP expression in response to the proteasome inhibitor bortezomib and observed the expected increase (Fig. 3b). In contrast to the suppression by 1-ABT seen with tioxazafen and wact-2 (Fig. 3b and 3f), 1-ABT enhances bortezomib activity (Figs. 3b, 3g and 3h), which suggests that P450s detoxify bortezomib in worms. As expected, both bortezomib and tioxazafen significantly upregulate GFP-tagged proteasome subunits (*p*<0.001) (Supplemental Fig. 6).

The above results indicate that bioactivated tioxazafen induces proteasomal stress. The root cause of this might either be global protein unfolding or inhibition of the proteasome itself. However, the sustained induction of *hsp-70* and *hsp-90* expression argues for the latter model (Bush *et al*. 1997). A second line of experimentation that can help resolve the two models is determining whether polyubiquitinated proteins accumulate in tioxazafen-treated worms. Generally, an intact proteasome can process polyubiquitinated proteins such that they fail to accumulate over a sustained period (Bence *et al*. 2001; Lehrbach and Ruvkun 2019; Lee and Goldberg 2022). We therefore invested whether polyubiquitinated proteins accumulate in tioxazafen-treated worms over a sustained period by exploiting a UIM2::ZsProSensor poly-ubiquitination reporter (Matilainen *et al*. 2013). ZsProSensor is a fluorescent protein (ZsGreen) that is fused to ornithine decarboxylase, which is degraded by the proteasome in a ubiquitin-independent manner (Momose *et al*. 2012). UIM2 is a ubiquitin-binding domain. Upon binding polyubiquitinated proteins, the UIM2::ZsProSensor is protected from being proteolytically degraded. Hence, the UIM2::ZsProSensor perdures only upon the accumulation of polyubiquitinated proteins, which occurs robustly upon proteasome inhibition. Indeed, tioxazafen and wact-2 induce UIM2::ZsProSensor accumulation in a P450-dependent manner (Fig. 3i-3o). The positive control, bortezomib, also induces the accumulation of the sensor (Figs. 3i, 3n, and 3o). Consistent with these results, we also found tioxazafen to cause accumulation of Ub[G76V]::GFP, which is normally degraded in a proteasome-dependent manner (Dantuma *et al*. 2000; Segref *et al*. 2011) (Supplemental Fig. 7). Together, these results support a model whereby bioactivated tioxazafen inhibits the proteasome.

### Bioactivated Tioxazafen Induces a Proteasome Bounce-Back Response in *C. elegans*

Analyses with our stress reporters suggest that the worm has a robust transcriptional response to bioactivated tioxazafen. To investigate further, we performed RNAseq experiments to identify the genes that are distinctly regulated in response to bioactivated tioxazafen. We collected synchronized first-stage larvae (L1s) and incubated them for 18 hours in either 7.5 μM tioxazafen (+/- 1 mM 1-ABT), 7.5 μM selectivin (+/- 1 mM 1-ABT), or solvent-only controls (+/- 1-ABT). Three biologically independent experiments were done for each sample.

To identify genes that are transcriptionally up-regulated in response to bioactivated tioxazafen, we first identified those whose transcription increases in response to tioxazafen relative to solvent-only control using a 2-fold-or-more cutoff at a false discovery rate (FDR) of *p*<1E-5. This comparison yielded 1396 genes whose transcription is up-regulated in response to either the parent tioxazafen molecule and/or any P450-dependent metabolites (Fig 4a; Datafile 1). We then identified genes whose transcription increases in response to tioxazafen + 1-ABT relative to solvent-only + 1-ABT using the same statistical criteria outlined above. This yielded 457 genes that are up-regulated in response to the tioxazafen parent molecule only. By subtracting the second list of genes from the first, we identified 1066 genes that are up-regulated in response to P450-bioactivated tioxazafen (Fig 4a; Datafile 1). We performed the same analysis with selectivin. A comparison of the two data sets yields 931 genes that are distinctly up-regulated in response to bioactivated tioxazafen (Fig. 4a). We repeated the identical analysis, but this time focusing on down regulated genes. We identified 717 genes that are down-regulated in response to bioactivated tioxazafen (Fig 4b; Datafile 1).

**Figure 4.**
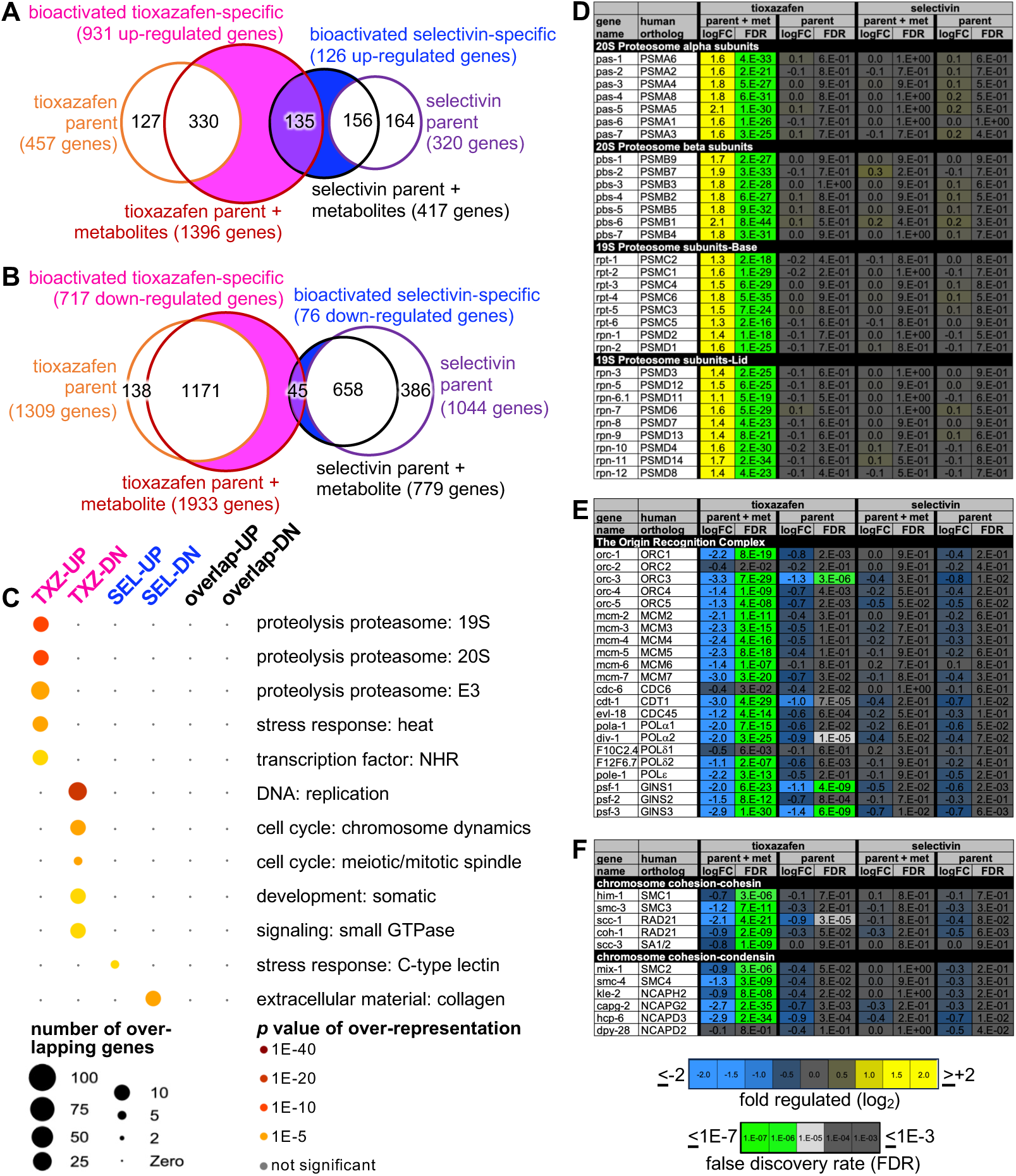
Bioactivated Tioxazafen Up-Regulates Proteasome Transcription Coupled to a Transcriptional Down-Regulation of Components Involved in the Cell Cycle. **A.** A Venn diagram showing the overlap of different categories of up-regulated genes (2-fold up-regulated or more with a false discovery rate of less than 1E-5; N=3 for each experimental subset). **B.** The same as A, but for down-regulated genes. **C.** Wormcat analysis of gene category enrichment. Only categories with a False-Discovery Rate-(FDR) corrected Fisher *p* value of less that 1E-5 are shown. TXZ, tioxazafen; SEL, selectivin. **D-F.** The log fold-change (LogFC) and FDR for the indicated categories (N=3, n=3 for each replicate).

We next asked which classes of genes are selectively impacted by bioactivated tioxazafen using the online Wormcat tool (Holdorf *et al*. 2020). We found that genes encoding proteasome subunits were significantly and distinctly up-regulated in response to bioactivated tioxazafen as a group (*p*<1E-9) (Fig. 4c and Datafile1) and individually (FDR<1E-11)(Fig. 4d). The up-regulation of proteasome subunit expression is the hallmark of the proteasome bounce-back transcriptional response that occurs upon proteasome disruption (Mannhaupt *et al*. 1999; Mitsiades *et al*. 2002; Lehrbach and Ruvkun 2016). By contrast, genes that are down-regulated selectively by bioactivated tioxazafen are involved in the cell cycle (Figs. 4c, 4e and 4f; Supplemental Fig. 8).

In *C. elegans*, the proteasome bounce-back transcriptional response is mediated by the NRF1 transcription factor ortholog SKN-1A (Lehrbach and Ruvkun 2016). In non-stressed conditions, the proteasome degrades SKN-1A. Upon proteasome disruption, SKN-1A is less efficiently degraded, accumulates, and promotes the transcription of proteasomal subunits. We therefore investigated whether bioactivated tioxazafen induces SKN-1A::GFP accumulation (Lehrbach and Ruvkun 2016), which we found to be true (*p*<0.001) (Fig. 5a-5n). The increase in SKN-1A::GFP is suppressed by 1-ABT, indicating that the effects are due to P450-bioactivated tioxazafen. By contrast, bortezomib does not induce obvious SKN-1A::GFP accumulation under these culture conditions (Fig. 5a-5n). Given that SKN-1A::GFP’s transcription is driven by the *rpl-28* promoter (Lehrbach and Ruvkun 2016), we examined our RNAseq data to examine whether the reporter’s unexpectedly high accumulation might be due to tioxazafen-driven *rpl-28* up-regulation. We found that *rpl-28* transcript abundance does not increase in response to tioxazafen (Datafile 1). Hence, bioactivated tioxazafen elicits a uniquely robust accumulation of SKN-1A protein, consistent with severe proteasomal disruption.

**Figure 5.**
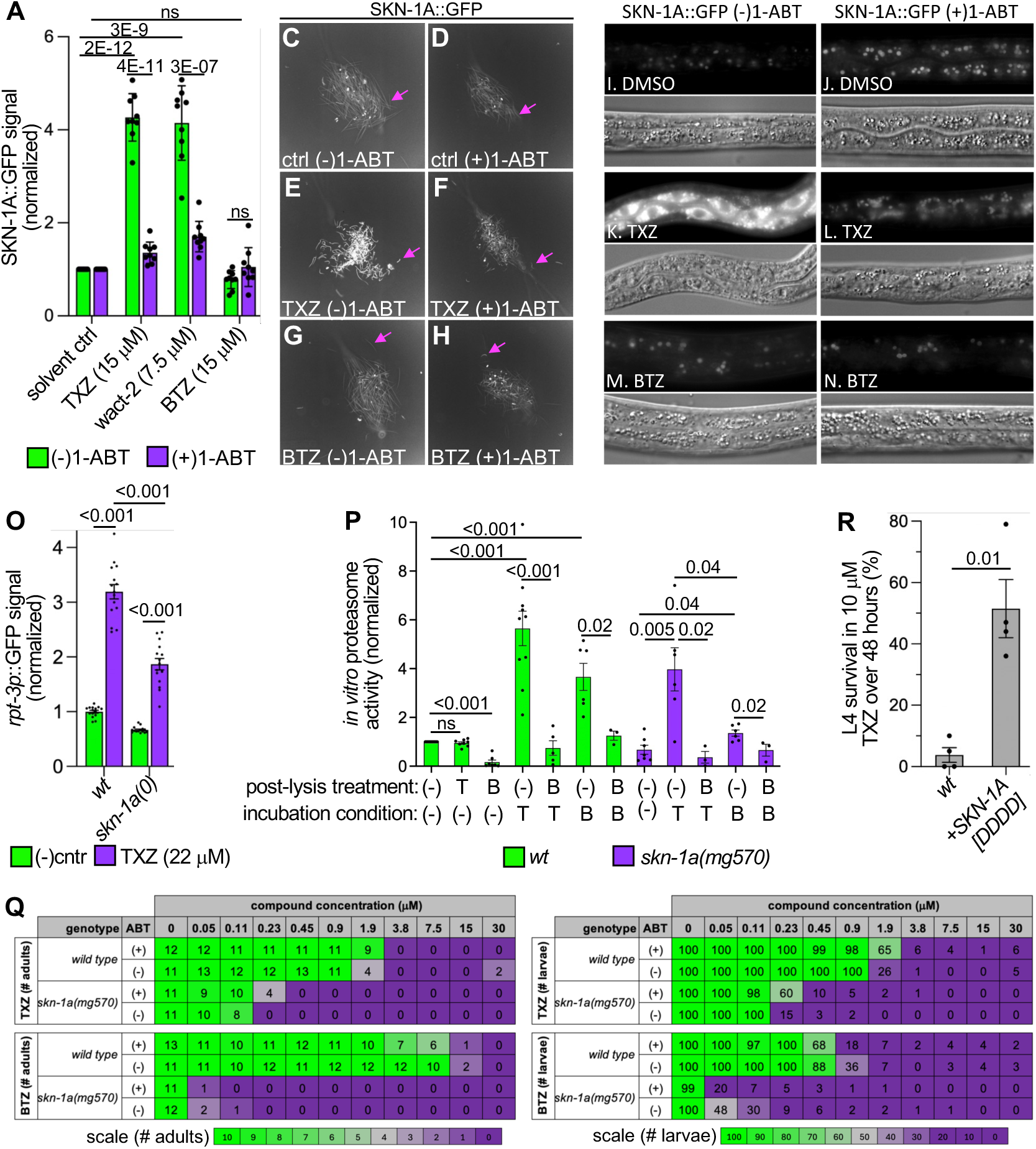
Tioxazafen Treatment Induces SKN-1A Accumulation, SKN-1A-Dependence, and Proteasome Inhibition. **A.** The accumulation of SKN-1A::GFP in the GR2198 strain in the indicated backgrounds. Nine independent trials (N=9) with at least 6 technical repeats per trial (n=6), with ∼150 worms per well in each technical repeat, was performed. Significance was determined using a two-tailed Student’s T-test. Standard deviation is shown. **C-H.** Examples of the SKN-1A::GFP fluorescence observed in the wells of selected experiments from ‘A’. The pink arrow highlights a single worm in each of the wells. **I-N.** Higher magnification images showing the accumulation of SKN-1A::GFP in animals incubated in the indicated compounds (15 μM) relative to controls. In A-N, worms were pre-incubated with 1-ABT or control for 24 hours, followed by incubation with either solvent-only, tioxazafen (TXZ) or bortezomib (BTZ). The mid posterior of a single animal is shown in each panel. With each pair of images, the top shows SKN-1A:GFP; the bottom shows a corresponding bright field image. **O.** Tioxazafen’s upregulation of the *rpt-3* reporter is only partially dependent on *skn-1a(mg570)*. L4 animals were exposed to 5 μM tioxazafen or vehicle control for 16 hours before imaging. Quantification of n=15 animals per condition is shown. Error bars show mean ±SD. Significance was determined by ordinary two-way ANOVA with Šidák’s multiple comparisons test (Note: multiple comparisons test applied to include additional comparisons as shown in fig S8). **P.** *In vitro* proteasome activity for the indicated genotype and treatment. Animals were incubated in either 1% solvent-only (-), 7.5 μM tioxazafen (T) or 15 μM bortezomib (B) for 18 hours, rinsed, lysed and the lysate was then either treated with solvent only (-), 7.5 μM tioxazafen (T) or 15 μM bortezomib (B) and immediately assayed for proteasome activity. N≥3, n=1. A non-parametric Mann-Whitney U test was used to assess significance. S.E.M. is shown. **R.** The survival of L4s of the indicated genotype placed on plates with 10 μM tioxazafen for 48 hours. (N=4, n=20 animals per replicate). Error bars show mean ±SD. An unpaired Welch’s T Test was used to calculate significance. **Q.** A dose-response for the indicated molecule. The number of adults (left) and larvae (right) were counted 6 days after wells were seeded with ∼20 L1 animals; the averages are reported. Note counting larvae are capped at 100; N=6 independent repeats, with n= 3 or 4 technical repeats each. In all panels, TXZ, tioxazafen; BTZ, bortezomib. In experiments shown in A-O, all samples were pre-incubated with either DMSO ((-)1-ABT) or 1 mM 1-ABT ((+) 1-ABT) for 24 hours.

Finally, we asked whether tioxazafen-induced proteasome upregulation is dependent on SKN-1A and found partial dependence (*p*<0.001) (Fig 5o; Supplemental Fig. 9). This contrasts with the proteasomal upregulation that is induced by bortezomib, which is entirely dependent on SKN-1A (Lehrbach and Ruvkun 2016; Lehrbach and Ruvkun 2019). Hence, bioactivated tioxazafen induces proteasomal up-regulation through both SKN-1A-dependent and independent mechanisms, which distinguishes it from known proteasomal disruptors.

### The Bounce-Back Response Induced by Bioactivated Tioxazafen is Evident *in vitro*

To investigate how bioactivated tioxazafen alters proteasome activity more directly, we measured proteasome activity in *C. elegans* lysates using a fluorogenic substrate that releases a fluorescent signal upon proteasome-specific cleavage (see methods). We first asked whether the parent tioxazafen molecule can suppress proteasome activity when added post-lysis of wild type control worms. In contrast to the positive control bortezomib, unmetabolized tioxazafen cannot inhibit proteasome activity (*p*>0.05) (Fig. 5p). As expected, this indicates that without P450-mediated bioactivation, the parent molecule cannot disrupt proteasome activity.

In parallel, we asked how proteasome activity is altered in worms incubated for 18 hours with tioxazafen or the bortezomib positive control. In both samples, we observed a significant increase in proteasome activity post-lysis (*p*<0.001)( Fig. 5p). With both samples, the increased proteasome activity can be suppressed by adding bortezomib post-lysis (*p*<0.05). Bortezomib is a reversible proteasome inhibitor (Williamson *et al*. 2006). Because of the bounce-back response, bortezomib incubation in worms leads to increased proteasome production. Upon lysis however, the reversable inhibitor is diluted and leaves evidence of the bounce-back response in its wake. Like bortezomib, we infer that bioactivated tioxazafen is inhibiting the proteasome reversibly.

In parallel, we also asked whether the bounce-back response that is evident in the *in vitro* assay is SKN-1A-dependent. *skn-1a(0)* mutants incubated in bortezomib show a small increase in proteasome activity relative to the *skn-1a(0)* control (*p*=0.01) and this increase is suppressed by post-lysis incubation in bortezomib (*p*=0.02)( Fig. 5p). This suggests that even bortezomib can induce a modest SKN-1A-independent bounce-back response. By contrast, *skn-1a(0)* mutants incubated in tioxazafen have a large increase in proteasome activity relative to the *skn-1a(0)* control (*p*=0.005), which is also suppressible by post-lysis addition of bortezomib (*p*=0.02) (Fig. 5p). These results are consistent with the transgenic analyses presented above that shows that the tioxazafen-induced bounce-back response is only partially dependent on SKN-1A.

### Bioactivated Tioxazafen Kills via Proteotoxic Stress

The proteasome is essential in *C. elegans* (Takahashi *et al*. 2002). If tioxazafen-induced lethality is caused by proteasomal stress, then worms that have a compromised bounce-back response (and therefore have fewer proteasome subunits under proteasomal stress) should be hypersensitive to tioxazafen. To test this, we asked whether *skn-1a(0)* null mutants are hypersensitive to tioxazafen and found that they are (*p*<1E-5), but less so than the mutant’s hypersensitivity to bortezomib (Fig. 5q). Again, this is consistent with the above results that show that the bounce-back response induced by tioxazafen is only partially dependent on SKN-1A. *skn-1a(0)*’s hypersensitivity to tioxazafen is at least partially suppressed by 1-ABT, consistent with proteasomal stress induced by a P450-dependent metabolite (*p*<0.05)(Fig. 5q). We also asked whether the converse is true. That is, whether animals with upregulated SKN-1A activity via the *skn-1a*[DDDD] transgene (Lehrbach and Ruvkun 2019) is sufficient to suppress tioxazafen-induced lethality. Indeed, we found that the *skn-1a*[DDDD] transgenic animals survived tioxazafen significantly better than the wild type controls (*p*<0.01; Fig. 5r). Together, the results show that proteasomal stress is a major contributor to how bioactivated tioxazafen kills animals.

### Humans Cytochrome P450s Bioactivate Tioxazafen

Given that some humans reportedly develop rashes after contacting tioxazafen-coated seeds, we wondered whether any human P450s are capable of toxifying tioxazafen. We therefore employed the PIXY assay to survey the 23 human P450s from the major drug metabolizing families (1-3) for their ability to bioactivate tioxazafen. We found that yeast expressing CYP1A1, CYP1B1, and to a lesser extent, CYP2C19, grow significantly slower in the presence of tioxazafen (*p*<1E-4) (Fig. 6a). A detailed dose-response analysis confirmed the CYP1A1 and CYP1B1 results (Fig. 1d). LC-MS analysis showed that many of the same tioxazafen metabolites are produced in these cells compared to worms and yeast expressing nematode P450s (Supplemental Fig. 10). We conclude that human CYP1A1 and CYP1B1 can toxify tioxazafen.

**Figure 6.**
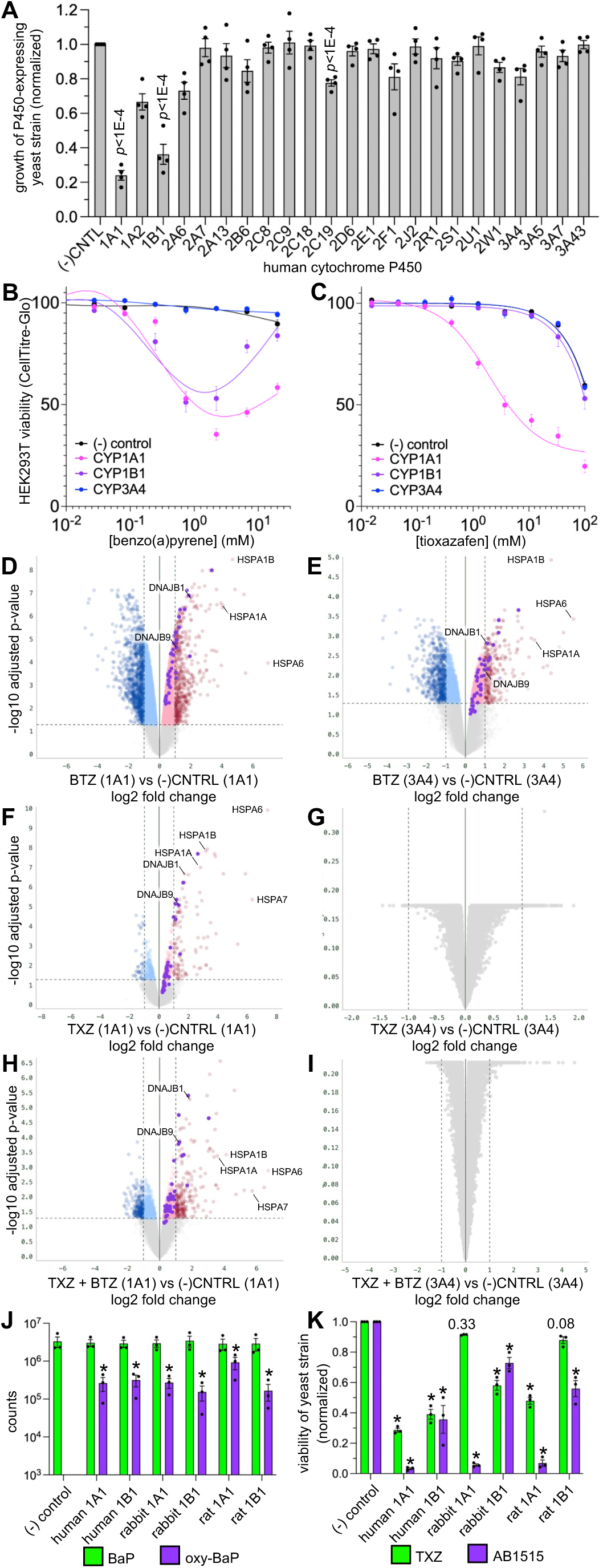
Tioxazafen is Bioactivated by human cytochrome CYP1A1 and CYP1B1. **A.** Viability of yeast strains expressing the indicated human P450 and incubated in 100 μM tioxazafen for 48 hours in the PIXY assay while measuring OD600 every 30 minutes. Area under the curve (AUC) was measured and normalized to the AUC of the empty-vector (-) control strain. A one-tailed Students T-Test was used to calculate significance relative to the (-) control with significant *p* values indicated. N≥3, n=2. **B-C.** A dose-response analysis of the ability of BaP and tioxazafen, respectively, to be toxified by the indicated P450 or negative control stably transduced HEK293T cells. The LD50 for tioxazafen with CYP1A1 and 1B1 is 2.2 mM and 9.8 mM, respectively. BaP follows a bi-phasic pattern of lethality in cells over-expressing CYP1A1 and CYP1B1. **D-G.** Bulk RNAseq analyses of CYP1A1-expressing HEK293T cells (left) or CYP3A4-expressing control HEK293T cells (right) with either 10 nM bortezomib (D-E) or 10 μM tioxazafen (F-G) for 24 hours. **H.** Combining the datasets from D and F reveals common genes that are significantly upregulated in response to BTZ and TXZ in cells over-expressing CYP1A1. **I.** Combining the datasets from E and G reveals no genes that are commonly upregulated in response to BTZ and TXZ in cells over-expressing CYP3A4. In D-I, datapoints highlighted in purple are Hallmark Human 2025 genes that belong to the unfolded protein response group. N=3. **J.** LC-MS analyses of the ability of the indicated yeast-expressed P450s to oxidize 100 μM benzo[a]pyrene (BaP) (parent exact [H+] mass 253.1012; oxidized mass 269.0961). Statistical differences relative to the negative control was done on Log_10_-transformed data using a student’s t-test. Asterisks = *p*<0.01, N=3, n=2. **K.** Yeast viability assays as described above with the indicated compound incubated at 100 μM. Two-way ANOVA was used to calculate significance relative to the (-) control with the asterisks indicating *p*<0.0001 with *p* values otherwise indicated. N=3, n=2. In all graphs, S.E.M. is shown.

We next asked whether expression of these P450s in HEK293T human-derived cells can also toxify tioxazafen. We integrated CMV-driven human CYP1A1, CYP1B1, or CYP3A4 (as a negative control) into HEK293T via lentivirus at a high MOI and selected for successful integrations with puromycin. As a control, we tested whether CYP1A1 and 1B1-expressing cells could toxify benzo(a)pyrene (Quan *et al*. 1994), and found that they could in an expected biphasic pattern (Spink *et al*. 2002; Genies *et al*. 2013) while control cell lines failed to do so (Fig. 6b). We then tested which P450s could toxify tioxazafen and found that expression of CYP1A1, but not CYP1B1, kills HEK293T cells in a tioxazafen-dependent manner with an LD50 of 2.1 μM (Fig. 6c).

We investigated whether the HEK293T cells respond to bioactivated tioxazafen in a similar manner to the positive control bortezomib by measuring transcriptional changes via bulk RNAseq. We conducted pilot tests with different concentrations of compounds and incubation times, all of which reveal similar trends (Supplemental Fig. 11; Datafile 1). For example, treatment of CYP1A1-expressing cells with 10 nM bortezomib for 24 hours revealed significantly up-regulated functional groups of genes that include the unfolded protein response, apoptosis, TNFA signaling via NFKB, and IL6 JAK STAT3 signaling (enrichment >1.5 fold; *p*<0.001), none of which is dependent of CYP1A1 expression (Fig. 6d-e; Datafile 1). As expected for proteasomal inhibition and the subsequent accumulation of unfolded proteins (Bush *et al*. 1997), HSP40 (DNAJB family) and HSP70 (HSPA family) expression is induced by bortezomib in a P450-independent manner (Fig. 6d-e; Datafile 1). A classic proteasomal bounce-back response was not evident but this is not unexpected given that proteasomal inhibitors can have narrow activity windows in which the bounce-back response is evident, depending on the cell line (Sha and Goldberg 2014). Treatment of CYP1A1-expressing cells with 10 μM tioxazafen for 24 hours resulted in similar groups of genes being upregulated, including all those listed above for bortezomib treatment (*p*<0.001) (Fig. 6f; Datafile 1). This transcriptional response to tioxazafen was entirely dependent on CYP1A1 (Fig. 6g; Datafile 1). Further highlighting the similarity of the cells’ response to bortezomib and bioactivated tioxazafen, combining the expression data from the two treatments in CYP1A1-expressing cells yields many of the same up-regulated genes (Fig. 6h), while doing the same with CYP3A4-expressing cells data yields no significantly regulated genes (Fig. 6i). Hence, bioactivated tioxazafen elicits a similar stress response in human cells as a known proteasome inhibitor.

Tioxazafen’s safety was evaluated using multiple animal models in pre-market regulatory tests, yet robust skin reactions were not evident (Health Canada 2017). At most, dermal assessment in rabbits was classified as slightly irritating. There is significant divergence of animal model CYP1A1 and 1B1 sequence from humans of up to 24% (Supplemental Fig. 12). We therefore wondered to what extent could CYP1A1 and CYP1B1 from mammalian models toxify tioxazafen, which we tested in our yeast-expression assay. We first ensured that the yeast-expressed animal model P450s were functional by assaying their ability to metabolize Benzo-a-pyrene (BaP), a known substrate of CYP1A1 and 1B1 (Bauer *et al*. 1995; Kim *et al*. 1998), and found that they can (Fig 6j). The P450s were also capable of toxifying a small molecule probe that we use in the lab as a toxified positive control called AB1515 (Fig 6k). However, rabbit CYP1A1 and rat CYP1B1 cannot robustly toxify tioxazafen (*p*>0.05; Fig. 6k). These data provide a reasonable explanation for why pre-market tests might have failed to reveal safety liabilities of tioxazafen.

## Discussion

We previously identified tioxazafen and several analogs in our screen for novel nematicides (Burns *et al*. 2015). These molecules piqued our interest because they were also revealed in our subsequent screen for molecules whose lethality is P450-dependent (Knox *et al*. 2024). Here, we provided multiple lines of evidence that show that toxified tioxazafen disrupts the proteasome (Fig. 7). P450-toxified tioxazafen results in the obvious accumulation of poly-ubiquitinated proteins, which is a tell-tale sign of proteasome inhibition (Menendez-Benito *et al*. 2005; Obeng *et al*. 2006; Higgins *et al*. 2015; Lehrbach and Ruvkun 2016). Consistent with this, toxified tioxazafen induces a dramatic increase in SKN-1A accumulation and a compensatory increase proteasome subunit abundance. Upon dilution of inhibitory factors upon cell lysis, the increased abundance of functional proteasomes becomes evident. Worms lacking SKN-1A are hypersensitive to tioxazafen. Conversely, worms expressing hyperactive SKN-1A resist tioxazafen’s lethality. Hence, proteasome inhibition is physiologically relevant to tioxazafen’s lethality. Notably, the P450-bioactivated nematicide selectivin does not induce a proteasomal bounce-back transcriptional response or interact with the proteasome in yeast, indicating that tioxazafen is distinct among pro-nematicides.

**Figure 7.**
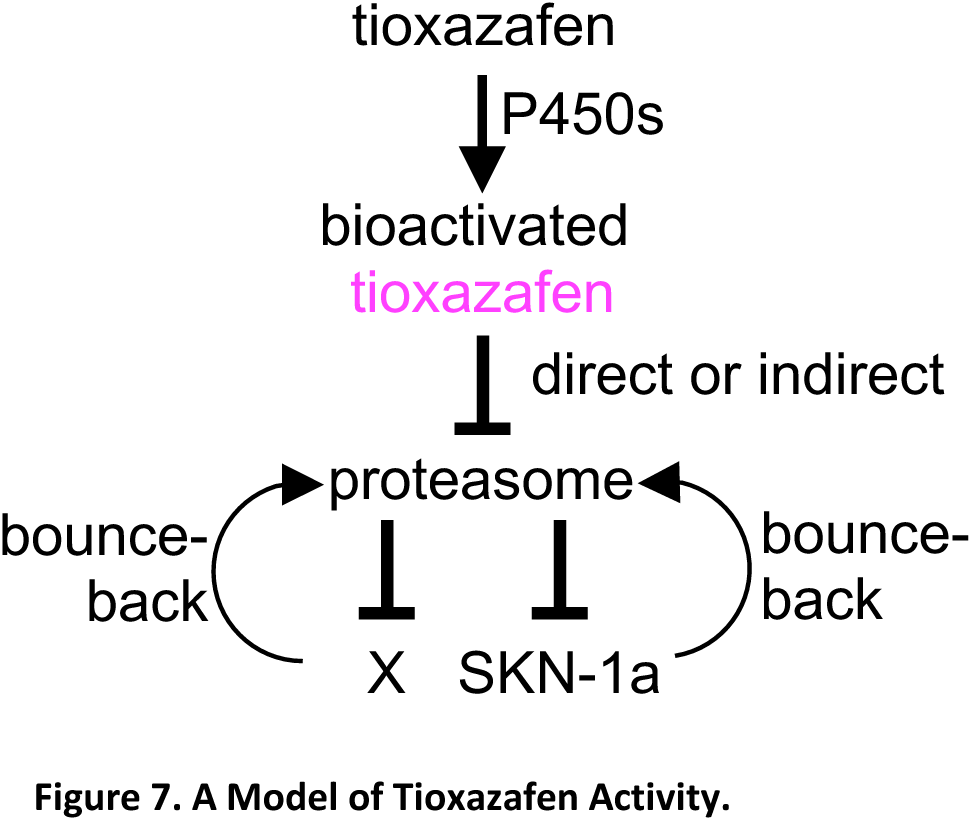
A Model of Tioxazafen Activity.

By multiple measures, tioxazafen has a distinct mode of action compared to selectivin and cyprocide. First, both selectivin and cyprocide are metabolized into electrophilic products that readily conjugate to low molecular weight thiols (LMWTs) in *C. elegans*. By contrast, no LMWTs conjugates of tioxazafen are observed in worms. It is likely that tioxazafen’s thiophene ring is oxidized by the P450s into an unstable reactive epoxide, which is common with structures of this type (Guengerich 2003). Given the absence of LMWT conjugates in worms, epoxidated tioxazafen must be rapidly consumed by some molecule(s) in the cell such that LMWTs fail to have an opportunity to react with it. Second, the pattern of global genetic interactions of bioactivated tioxazafen is distinct from that of bioactivated selectivin; the former genetically interacts with mutant proteasomal components while the latter interacts with mutant protein export machinery. Finally, both the large-scale stress reporter survey and RNAseq data show that tioxazafen elicits a distinct stress response in the worm compared to selectivin; the tioxazafen elicits a proteasomal bounce-back and an unfolded protein response, while selectivin fails to elicit a meaningful stress response at the concentrations that we have tested. Hence, bioactivated tioxazafen is unique in its mode of action of the pro-nematicides so far discovered.

The mechanism by which toxified tioxazafen inhibits the proteasome remains unclear. Given that inhibition is reversible upon cell lysis, it is unlikely that a reactive tioxazafen metabolite is covalently disrupting the proteasome. Instead, we speculate that a reactive tioxazafen metabolite binds a non-proteasomal component in the cell such as a peptide that in turn inhibits the proteasome. Identifying the tioxazafen-based inhibitor could prove to be a useful addition to our arsenal of proteasome disruptors.

Tioxazafen’s effect on the proteasome and its regulation is distinct from that of bortezomib in several respects. First, the effect on SKN-1A levels is distinct. SKN-1A::GFP accumulates dramatically in tioxazafen-treated worms, even though under the same culture conditions, we could not detect BTZ-triggered SKN-1A accumulation. Importantly, *skn-1a*(*0*) mutants are hypersensitive to both drugs, indicating that although present at different levels, SKN-1A is functional in both cases. Second, the effect on SKN-1A localization appears distinct. A large fraction of the accumulated SKN-1A::GFP in toixazafen-exposed animals is present in the cytosol. By contrast, the same SKN-1A reporter accumulates predominantly in the nucleus of following proteasome RNAi (Lehrbach and Ruvkun 2016) or inhibition by BTZ (Lehrbach and Ruvkun 2019). Third, tioxazafen causes upregulation of proteasome abundance in part via a SKN-1A-independent mechanism, whereas the effect of bortezomib is entirely SKN-1A-dependent. Consistent with this, the *skn-1a(0)* mutant’s hypersensitivity to tioxazafen is less than its hypersensitivity to bortezomib. Finally, toxified tioxazafen coincidentally downregulates the expression of cell cycle machinery whereas other proteasome inhibitors disrupt the cell cycle largely through post-transcriptional mechanisms (Ishii *et al*. 2006; Chen *et al*. 2011) and generally promote transcription (Yang *et al*. 2006; Kinyamu *et al*. 2020). Whether tioxazafen has this effect through the indirect modulation of cell cycle regulators via proteasome disruption (Pagano *et al*. 1995; Clurman *et al*. 1996; Tokumoto *et al*. 1997) or by some other mechanism is unknown. The proteasome is a complex machine, and it is unlikely that all modes of its inhibition have been discovered. It may be that a single tioxazafen-derived metabolite and/or conjugate is responsible for inhibiting the proteasome in a unique way that leads to the cell’s distinctive response. Alternatively, it may be that multiple tioxazafen metabolites and/or conjugates are collectively responsible for the distinctive effects of P450-bioactivated tioxazafen. In either case, further investigating how toxified tioxazafen exerts its inhibitory effects on the proteasome will likely yield novel insight.

Multiple experiments indicate that the *C. elegans* bounce-back response to bioactivated tioxazafen is only partially dependent on SKN-1A. In mammals, there are several other mechanisms by which the proteasome can be upregulated. For example, NRF2 can upregulate the proteasome in response to oxidative stress (Kwak *et al*. 2003; Pickering *et al*. 2012), while NRF3 is necessary to maintain proteasomal gene expression in at least some contexts (Waku *et al*. 2020). FOXO1 and FOXO4 transcription factors have also been shown to regulate proteasome subunit expression (Liu *et al*. 2017; Kapetanou *et al*. 2021). Additional pathways have been implicated (Vilchez *et al*. 2012; Kamber Kaya and Radhakrishnan 2021). Determining whether the orthologs of any of these proteins act in concert with SKN-1A to mediate the worm’s bounce back response to toxified tioxazafen awaits further investigation.

As introduced above, tioxazafen commercialisation efforts were halted because workers reportedly developed skin rashes after handling them. Our survey of human cytochrome P450s belonging to families 1-3 revealed that CYP1A1 and 1B1 toxify tioxazafen when expressed in yeast. We observe similar effects for CYP1A1 when expressed in human-derived HEK293T cells. In single cell sequencing databases, CYP1A1 and CYP1B1 are the most highly expressed P450s in skin (Abdulla *et al*. 2025), while other studies show that CYP1B1 and CYP2E1 are the most highly expressed (Chen *et al*. 2024). Regardless, it is CYP1A1 that is most highly induced by external factors such as ultraviolet-B light (Katiyar *et al*. 2000) and polycyclic aromatic hydrocarbon pollutants and pro-carcinogens (Von Koschembahr *et al*. 2018), which may be consistent with the idea of factory and farm workers being especially sensitive to the pro-nematicide.

Through our investigation, we have established yeast-based assays for the ability of nematode and human P450s to toxify what was an otherwise effective nematicide (Slomczynska *et al*. 2015). It may be possible to exploit these assays to develop tioxazafen analogs that will retain nematicidal properties while avoiding the toxic reactions in humans to yield a useful and safe agent to control plant parasitic nematodes globally.

## Supporting information

Supplemental Display Items

Datafile 1

## Acknowledgements

We thank Anchal Aggarwal for preliminary technical work; Igor Stagljar and Jamie Snider for training and use of their CLARIOstar fluorimeter; Jed Goldstone as a member of the International Committee on Cytochrome P450 Nomenclature for help with nomenclature. Several *C. elegans* strains were provided by the CGC, which is funded by NIH Office of Research Infrastructure Programs (P40 OD010440). We thank Stephen Bell and James De Voss for insights into potential tioxazafen reaction mechanisms.

## METHODS

### *C. elegans* Strains and Culture

*C. elegans* strains were maintained using standard culture practices (Kwok *et al*. 2006). All experiments were carried out at 20°C unless otherwise stated. The following strains were obtained from the *C. elegans* Genetics Centre (University of Minnesota) or our own collections: YD90 *vha-6p*::UIM2-ZsProSensor, N2 (wildtype), CL2166 *(dvIs19 [(pAF15) gst-4p::GFP::NLS] III),* CF1553 (*muIs84* [*sod-3p::GFP* + *rol-6*(su1006)]), AU133 *(agIs17 [irg-1p::GFP + myo-2p::mCherry] IV),* IG274 *(frIs7 [nlp-29p::GFP + col-12p::DsRed] IV),* IG1295 (*frIs21*[*cnc-2p::GFP*]), VP198 *(kbIs5[gpdh-1p::GFP + rol-6(su1006)])*, SJ4005 *(zcIs4 [hsp-4::GFP] V),* SJ4100 *(zcIs13 [hsp-6::GFP])*, TJ375 *(gpIs1[hsp-16.2p::GFP]),* GR2183 (*mgIs72 [rpt-3p::GFP + dpy-5(+)*] *II*), GR2197 *mgIs72[rpt-3p::gfp] II; skn-1a(mg570) IV;* GR2198 *mgTi1 rpl-28p::skn-1a::GFP::tbb-2 3’UTR,* GR2211 *mgIs72[rpt-3p::gfp] II; ddi-1(mg571) IV*; GR2215 *mgIs72[rpt-3p::gfp] II; sel-1(mg547) V*; GR2236 *png-1(ok1654) I; mgIs72[rpt-3p::gfp] II;* GR2245 *skn-1a(mg570);* GR3090 *mgIs77[rpl-28p::ub(G76V)::GFP] V*; GR3094 *skn-1(mg570) IV; mgIs77[rpl-28p::ub(G76V)::GFP] V*; NJL4774 *pbs-4(nic1600[pbs-4::sfGFP]) I*; NJL3930 *unc-119(ed3) III; mgTi20[rpl-28p::SKN-1t(DDDD)]*; RB1613 *cyp-35A5(ok1985)*(a 807 bp deletion); WDC4 *rpn-8(ana4[gfp::rpn-8]) I*. QV225 *(skn-1(zj15) IV)* was provided by Keith Choe; *rmIs8 [hsp-70::GFP])* and AM799 (*rmIs317[daf-21p::GFP]*) were provided by Richard Morimoto; IG1295 (*frIs21*[*cnc-2p::GFP*]) was a kind gift from Jonathan Ewbank; the YD90 *vha-6p*::UIM2-ZsProSensor strain was a kind gift from Carina Holmberg (University of Helsinki). *cyp-35D1(ean222)*, a deletion allele, was a kind gift from JB Collins and Erik Andersen, and VC40639 *cyp-33c2(gk738011)* (a C>T substitution, which translates to a premature Q276X stop) was from the million mutation project (Thompson *et al*. 2013).

To synchronize *C. elegans* larvae, we use a previously described protocol (Burns *et al*. 2006). Briefly, animals are cultured on 10 cm nematode growth medium (NGM) solid plates seeded with *Escherichia coli* HB101 at 20°C until near confluence with many adults and embryos visible. Animals are washed off with M9 buffer and resuspended in 1.5 ml M9 buffer in a 15-ml conical tube with no more than 500 ml of packed worms in the 1.5 ml suspension. 4.5 ml alkaline bleach solution (10% NaOCl, 1M NaOH, and ddH2O) is then added to the worm suspension. The tube is then inverted immediately and twice, then once every 30 seconds for 4.5 min to prevent the worms from collecting at the bottom of the tube. The tube is then vortexed for 10 seconds. 6 ml of M9 buffer is then added to the digest and inverted 2-3 times. The tube is then immediately centrifuged for 2 min at 796g (2,000 r.p.m. in a benchtop Eppendorf 5810 R centrifuge). The supernatant is the aspirated and the worms are resuspended in 10 ml of M9 buffer. The tube is then shaken manually until the pellet is resuspended and the process is repeated two more times. The resulting embryos are then resuspended in 10 mLs of M9 solution in 15 mL Falcon tubes and placed on a rotator overnight at 20°C to yield synchronized L1-stage larvae.

### *emb-8(RNAi)* Dose Response

*E. coli* HT115(DE3) containing an RNAi feeding vector expressing dsRNA targeting the *emb-8* gene or an empty RNAi feeding vector (L4440) were grown overnight in LB with 100 µg/mL carbenicillin at 25°C with no shaking until the culture was in mid-log phase (OD600 ∼0.6). The culture was induced with 1 mM IPTG and grown at 37°C with shaking at 200 rpm for 1 hour. The bacteria were pelleted and concentrated ten-fold with liquid NGM containing 1mM IPTG and 100 µg/mL carbenicillin. This bacterial suspension was dispensed into the wells of a flat-bottomed 96-well culture plate (40 µL per well). *C. elegans* temperature sensitive mutant strain MJ69 *emb-8(hc69)* and wild-type strain N2 synchronized L1s were obtained from an embryo preparation that was left to hatch at 15°C overnight. Approximately 20 N2 L1s were plated in 10 µL of M9 per well containing L4440 empty vector control bacteria (“wild type” condition). Approximately 20 *emb-8* L1s were plated in 10 µL of M9 per well containing *emb-8* dsRNA expressing bacteria (“POR knockdown” condition). Worms were grown for 40 hours at the restrictive temperature of 25°C with shaking at 200 rpm at which point 120 µM FUDR was added to each well and plates were returned to the 25°C incubator for another 30 hours. At this 70 hour timepoint the compounds were added to the wells using a multichannel pipette to a final DMSO concentration of 1% v/v. After 48 hours of chemical exposure viability was quantified by observation under a dissection microscope, and the data is presented as the proportion of viable worms in each condition relative to the DMSO controls. Three biological replicates were completed with two technical replicates on each day and the proportion viable was averaged across the replicates.

### Mutant P450 Dose Response Assay

50-100 L1s of either WT, RB1613 *cyp-35A5(ok1985)*, *cyp-35D1(ean222)*, or VC40639 *cyp-33c2(gk738011)* in 20 µL of M9 buffer were added to each well of a 96-well plate (Sarstedt, Nümbrecht, Germany) containing 80 µL of culture media (80 uL of *E. coli* HB101 bacteria (OD_600_ = 2.0-2.2) and liquid NGM). Using a drug pinning tool/replicator (V & P Scientific Inc., San Diego, California, U.S.A.), 0.3 µL of 1% DMSO (solvent control) or increasing concentrations (0.78-25 µM) of Tioxazafen or wact-2 was added to each well in triplicate. The plate was sealed with parafilm and placed within a closed plastic container containing wet paper towels. The plate was incubated for 3 days at 20°C with horizontal shaking at 2,000 RPM (New Brunswick Scientific, New Jersey, U.S.A.) in the dark. On the third day, worms were briefly spun (5 seconds at 1,000 RPM) (Sorvall X Pro Series, Thermo Scientific, Massachusetts, U.S.A.). Worm motility per well was assessed by gently dropping the plate from a height of ∼1 cm from the dissection microscope stage inspecting for brief/sustained thrashing of worms (motile) or lack of movement. The experiment was repeated at least 3-5 times. Statistical analyses were performed on GraphPad Prism 10 (Boston, MA 02110, USA) using non-parametric Mann-Whitney U-test (*p*<0.05).

### Worm Sample Preparation for LCMS Analyses

65,000 synchronized L1s were added to a sterile Eppendorf tube in a total volume of 500 µL M9 containing 1% DMSO (solvent control) or 100 µM Tioxazafen (in DMSO), in technical duplicates and incubated on a nutating mixer (VWR, Pennsylvania, U.S.A.) for 24 hours in the dark. After the incubation, worms were centrifuged at 6,000 RPM for 1 minute (Centrifuge 5417C, Hamburg, Germany), then transferred by low-retention tips to 96-well plates with filter bottom (Pall corporation, New York, U.S.A.). The plate was fitted on top of a 96-well chamber (Pall corporation, New York, U.S.A.) to hold aspirated buffer. The buffer was aspirated off each well and collected at the bottom plate and transferred to sterile Eppendorf tube. To each pellet on the top wells was added 50 µL double-distilled water, which was then carefully transferred by low-retention tips (the tip cut to about 2 mm with sterilized scissors) to sterile cryovials. Both the buffer and worm pellet for each sample was immediately stored at -80°C for extraction and LCMS analysis. The experiment was repeated at least three times. Statistical analyses were performed on GraphPad Prism 10 (Boston, MA 02110, USA) using non-parametric Mann-Whitney U-test (p<0.05).

### Yeast Sample Preparation for LCMS Analyses

Yeast strains harbouring plasmids capable of expressing P450s or the empty vector control with the relevant cofactor(s) were grown from a single colony overnight at 30 _o_C in 5 ml SD-URA+2% galactose to induce P450 and redox partner expression. The next morning, the OD600 of each culture was measured and concentrated to an OD600 of 10 in water. 0.5 mL aliquot of culture were treated with either 100 μM tioxazafen, 100 μM wact-2, 100 μM B[a]P or DMSO for 4 hours at 30 _o_C with constant mixing in a rotary incubator. Cells and buffer were then separated with AcroPrep™ Advance 96-well 0.45 µm WWPTFE filter plate (Pall Corporation, P/N 8684)using a vacuum manifold (Pall Corporation, P/N 5017).

Buffers were dried down at 60 _o_C using vacuum concentrator (Eppendorf, P/N 022820109) at the V-AQ setting (a pre-set mode for drying aqueous-based solutions) with no additional extraction steps.

Cells were lysed mechanically by adding 100 µL of 0.5 mm glass beads (BioSpec, P/N 11079105) and 800 µL of MeOH:BuOH 50:50 mixture with subsequent bead beating using Bead Ruptor homogenizer (Omni International, P/N 19-070). The homogenization protocol consisted of 5 cycles, each including 1 minute of bead beating at maximum speed followed by a 30-second dwell period. Following a spin at 14,000 rpm for 10 minutes to remove cell debris, the supernatant was transferred to a Safe-Lock Eppendorf tube (Eppendorf, P/N 022363204) and dried down at 60 _o_C using the Eppendorf concentrator at the V-AL setting (a pre-set mode for drying alcohol-based solutions). Both buffers and lysates were resuspended in 100 µL 50:50 ACN:H₂O, vortexed briefly, and sonicated for 15 minutes using the VWR ultrasonic cleanser (P/N97043-940) to ensure thorough resuspension. The resuspended samples were centrifuged at 14,000 rpm for 10 minutes, and 25 µL of the supernatant was transferred to polypropylene inserts (Agilent, P/N 5182-0549) in amber vials (Agilent, P/N 5182-0716).

### Liquid Chromatography Coupled Mass Spectrometry Analysis

An Agilent 1260 Infinity II with 6545 LC/QTOF mass spectrometer was used to analyze samples in positive ionization mode with Dual AJS electrospray ionization (ESI) equipped with Agilent ZORBAX Eclipse Plus C18 column (2.1×50mm, 1.8-μm particles) and ZORBAX Eclipse Plus C18 guard column (2.1×5mm, 1.8-μm particles). Liquid chromatography parameters used were: injection volume 5 μL with 5 μL needle wash with sample, autosampler chamber temperature 4 °C, column oven temperature 40 °C. Mass spectrometry parameters were: gas temperature 320 °C, drying gas flow 10 L/min, nebulizer 35 psi, sheath gas 350 °C at 11 liters per minute, VCap 3500 V, Nozzle voltage 1000 V, fragmentor 125 V, skimmer 65 V. The solvent gradient with the flow of 0.5 ml per minute started with 95% mobile phase A (Optima LC/MS H2O+0.1% Formic Acid, Fisher Chemical P/N LS118-4) and 5% mobile phase B (Optima LC/MS Acetonitrile+0.1% Formic Acid, Fisher Chemical P/N LS120-4), increased linearly to 50% B at 10 minutes, followed by an increase to 100% B at 27.5 minutes. The post-run time was two minutes (instrument conditioning at 95% mobile phase A). The raw MS scan was analyzed using Agilent MassHunter Qualitative Analysis 12.0 and Agilent MassHunter Profinder 10.0 to find differences in samples of different strains and conditions.

Tandem mass-spectrometry (MS/MS) fragmentation data for each mass of interest was obtained by doing targeted mass spectrometry acquisition with the same method parameters as described above. Constant collision energies of 10, 20, and 40 eV were applied for all molecules. The MS/MS data was then analysed and chemical structures predicted using MassHunter Qualitative Analysis 12.0 software, SIRIUS 6.3.4 software (Duhrkop *et al*. 2019) with fragmentation tree tool (Bocker and Duhrkop 2016), and CFM-ID online tool (Wang *et al*. 2022).

### Constructing Yeast Strains Expressing Human, C. elegans, Meloidogyne incognita, Rattus norvegicus, and Oryctolagus cuniculus P450s

The human P450 co-factors POR and cytochrome B5 (Accessions: P16435, P00167) (Amorosi *et al*. 2021), driven by the bi-directional *GAL1/GAL10* promoter (Johnston 1987) respectively, was integrated into the CAN1 locus of the yeast strain Y15771 MATa his3Δ leu2Δ ura3Δ pdr5Δ snq2Δ yor1Δ (Talkish *et al*. 2019) using homologous recombination (Ma *et al*. 1987). Briefly, sequence encoding the *3’ CAN1 seq*-*human POR-GAL1-10p-cytochrome b5-NATMX-5’ CAN1 seq* was inserted into a URA3 selectable, centromeric yeast expression vector by homologous recombination that also carries a constitutively expressed Cas9 and gRNA complementary to CAN1. The insert sequence is flanked by sequence complementary to the upstream and downstream regions of CAN1 to facilitate homologous recombination between the insert sequence and the CAN1 locus, which is stimulated by Cas9 cutting after plasmid transformation into yeast Y15771 by lithium acetate (Gietz and Schiestl 2007). Transformants that successfully took up the integrative plasmid were first selected for on SD-URA and allowed to grow for 48 hours at 30 °C. After 48 hours, colonies were replica-plated onto SD-Arg + canavanine, grown for another 48 hours at 30 °C, and subsequently replica-plated onto SD+all amino acids + NAT to select for colonies with a successful integration event. This new humanized strain is called RP5274 MATa his3Δ leu2Δ ura3Δ lys2Δ pdr5Δ snq2Δ yor1Δ can1::Human-POR-Human-B5. The RP5274 humanized base strain was also used for the rat (*Rattus norvegicus*) and rabbit (*Oryctolagus cuniculus*) P450 experiments. A similar protocol was used to generate the *C. elegans* base strain RP4400 MATa his3Δ leu2Δ ura3Δ pdr5Δ snq2Δ yor1Δ can1::gal-10/gal-1p::*C. elegans* EMB-8 HO::Barcode-HYGR. The *Meloidogyne incognita* P450 yeast experiments were done with the Y15771 base strain.

P450s of interest were expressed from the *GAL-1p* promotor in the centromeric low copy CEN6/ARS4 origin of replication ATCC p416 plasmid (Montalibet and Kennedy 2004). To build these, we obtained sequences from either WormBase (WS296) or Uniprot (for the mammalian P450s). The construction of the *M. incognita* P450-expressing plasmids was previously described (Knox *et al*. 2024). The codon-optimized the ORFs were synthesized by Twist Biosciences and placed into the SpeI and HindIII restriction sites of the ATCC p416. The resulting plasmids were transformed into the relevant base strains (described above) using lithium acetate methodology (Gietz and Schiestl 2007). All P450 sequences and resulting strain names are available in the DataFile 1.

### The PIXY Yeast Toxification Assay

The yeast strains described above were grown overnight at 30 °C in 5 ml SD-URA buffer (Bach and Lacroute 1972) + 2% raffinose. The next morning, the OD600 of each culture was measured and subsequently diluted back to an OD600 of 0.01 in SD-URA+2% galactose to induce P450 and redox partner expression. 0.2 ml of culture was aliquoted into the wells of a 96-well plate and treated with the relevant compounds or 1% solvent-only (DMSO) control. Plates were incubated for 48 hours at 30 °C, with OD600 measurements collected every 30 minutes. Area under the curve of the growth curves was calculated using Graphpad Prism 10.6.1 and normalised to DMSO and EV controls.

### Synthetic Genetic Array Analyses

We transformed the canonical SGA query strain Y7092 (*MATa can1Δ::STE2pr-Sp_his5*; *lyp1Δ; his3Δ1 leu2Δ0 ura3Δ0* LYS2) (Costanzo *et al*. 2016; Kuzmin *et al*. 2016) with either plasmid pPR2222, which is the empty vector control, plasmid pPR2125, expressing CYP-35C1, or plasmid pPR2126, expressing CYP-35D1, resulting in strains RP4222, RP4223 and RP4224, respectively. All plasmids are URA3-selectable of the CEN/ARS low-copy number type with inserts driven by the GAL1 promoter. With the resulting strains, we performed a dose-response analyses on the solid substrate SGA yeast growth media (Costanzo *et al*. 2016; Kuzmin *et al*. 2016) to determine the concentration of either tioxazafen or selectivin that reduces colony size growth of RP4223 and RP4224, respectively, by 30% relative to colony growth on solid substrate with solvent only control (1% DMSO). We performed the same dose response with the empty vector strain RP4222 to determine whether either compound affected the growth of yeast that do not express P450s on solid substrate. We found that 32 μM selectivin and 15 μM tioxazafen reduced growth of RP4223 and RP4224, respectively, by 30% but did not affect the growth of the RP4222 negative control strain.

For the SGA analyses, we used the complete nonessential gene deletion and essential gene temperature-sensitive (TS) allele arrays that are described elsewhere (Costanzo *et al*. 2016). Strains harboring the plasmids described above were crossed to the nonessential deletion and essential TS allele arrays as previously described (Costanzo *et al*. 2016; Kuzmin *et al*. 2016) with the following modifications. Nonessential deletion essential TS mutant strains carrying the plasmids described above were selected on medium supplemented with 2% glucose. Mutant arrays were subsequently grown on selective medium contain 2% raffinose followed by growth on selective medium containing with 2% galactose to induce CYP expression. Growth medium for mutant arrays expressing pPR2125(GAL1p-CYP-35C1) also contained 32 μM selectivin whereas growth medium for mutant arrays expressing pPR2126(GAL1p-CYP-35D1) was supplemented with 15 μM tioxazafen. Mutant arrays carrying the pPR2222 empty vector control were copied twice. One copy was grown in galactose-inducible medium containing 32 μM selectivin and the second copy was grown on galactose-inducible medium containing 15 μM tioxazafen. All resulting ‘doubles’ (i.e., the plasmid crossed into the mutant background) are pinned on to the solid substrate in four technical replicates. The entire screening pipeline was repeated three independent times.

To measure the outcome of the chemical-genetic interactions, we applied our previously described colony size scoring method (Baryshnikova *et al*. 2010; Wagih *et al*. 2013) to estimate fitness of mutant strains expressing CYP-35C1 or CYP-35D1 or the vector control in the presence of drugs. Colony size of SGA deletion and TS array mutant strains are measured from pictures of the plates taken 48 hours after incubation at 30°C. Nonessential gene deletion or essential gene TS alleles that enhanced or suppressed P450-dependent drug toxicity were identified by comparing fitness measurements for the same mutant strain expressing a P450 versus the vector control. P450-expressing mutants that exhibited reduced fitness relative to the vector control (i.e. negative scores) represented potential enhancers of P450-dependent drug toxicity whereas P450-expressing mutants that exhibited increased fitness relative to the vector control (i.e. positive scores) identified potential suppressors of P450-dependent drug toxicity.

Traditional GO enrichment analysis was performed as described elsewhere (Costanzo *et al*. 2016). Briefly Gene Ontology (GO) term enrichment was performed using a custom Fisher’s exact test pipeline built on GO annotations for *Saccharomyces cerevisiae* (SGD GAF, geneontology.org) (Ashburner *et al*. 2000) using a custom background gene list corresponding to all unique nonessential and essential genes that comprise the SGA mutant arrays (Costanzo *et al*. 2016). For each GO term, a one-tailed Fisher’s exact test assessed over-representation of the term in the study set relative to the custom gene background. The analysis was restricted to GO terms that comprised between 5 and 250 genes in the background. Odds ratios were calculated with a Haldane–Anscombe correction (+0.5) and resulting *p*-values were adjusted for multiple testing using the Benjamini–Hochberg false discovery rate procedure (adjusted *p* < 0.05 considered significant).

### *C. elegans* GFP Reporter Screens

Our liquid-based screening assay is an adaptation of a protocol used for multi-well liquid RNAi screens (Lehner *et al*. 2006). 160 mL of HB101 *E. coli* culture in LB broth was two-fold diluted in sterile nematode growth media (containing 3 mg/mL NaCl, 2.5 mg/mL Bacto-peptone, 5 µg/mL cholesterol, 1 mM CaCl2, 1 mM MgSO4, 25 mM KH2PO4). The resultant volume of media is sufficient to assay around 20 library plates. 90 µL of the HB101+NGM media was pipetted to each well of 96-well plates with a multichannel pipette. A 96-well pinning tool (V&P scientific) with a 300 (±20) nL volume slot was used to transfer chemicals from a stock library plates to the culture plates. With a multichannel pipette and siliconized tips, 20 µL of M9 solution that contained around 20 L4 worms were transferred to each well of the plates. The culture plates were then sealed with parafilm, covered with damp paper towel in Tupperware containers and placed in a 20 °C shaking incubator set at 200 rpm for 24 hours. Chemicals in our stock worm active (wactive) library are 10 mM in concentration and therefore the final screening concentrations of the library chemicals were around 30 µM (in 0.3% DMSO (v/v)).

20 synchronized L4 larval stage worms were dispensed in a total volume of 100 µl in each well of 96-well plates. Each well also contained 30 µM of each wactive. Screens were performed in duplicate. After 24 hours, GFP fluorescence intensity and pattern were evaluated with a Leica epifluorescence dissection microscope and recorded in a numerical scale of 0 to 3. A score of 0, 1, 2 or 3 represented no expression, faint expression, intermediate expression and strong GFP expression, respectively.

### *C. elegans* Fluorescent Reporter Analyses

A bacterial suspension in liquid NGM was prepared by concentrating a saturated overnight HB101 culture (OD_600_: 0.6-0.8). 40 μL of this bacterial suspension, containing either 1.25 mM 1-ABT or DMSO vehicle, was added to each well of a 96-well flat-bottom culture plate. Approximately 150 synchronized L1 YD90 *vha-6p*::UIM2-ZsProSensor, GR2198 *mgTi1 rpl-28p::skn-1a::GFP::tbb-2 3’UTR* and GR2183 *mgIs72 rpt-3p::GFP* worms were then added to each well in 10 µl of M9 buffer, achieving a final concentration of 1 mM 1-ABT. These L1 worms were incubated in the 1-ABT/DMSO-containing medium for 24 hours. Thereafter, 0.3 µl of tioxazafen, wact-2 or bortezomib was pinned from a multiwell plate into the 96-well plates containing worms using a 96-well pinning tool (V&P Scientific Inc). The culture plates were then incubated for an additional 24 hours at 20°C with shaking at 200 rpm. After this incubation, the plates were imaged using the Biotek Cytation 5 Imager (Agilent Inc). Fluorescence was quantified using ImageJ/Fiji software. With the GR2198 *mgTi1 rpl-28p::skn-1a::GFP::tbb-2 3’UTR* expedriment, we also photographed animals at higher magnification using a Leica DMRA 2 compound microscope with 400X magnification to examine subcellular distribution of the GFP fusion protein. Openlab software was used to capture images.

For the fluorescent analyses of of *rpt-3p*::GFP, PBS-4::GFP, GFP::RPN-8, and Ub[G76V]::GFP fusion proteins, NGM plates containing the desired concentration of tioxazafen (Aaron Chemicals, #AR01QXD0) or bortezomib (LC Laboratories, #B1408) were prepared by directly applying tioxazafen or bortezomib solution to NGM plates seeded with OP50 and leaving the plates to dry for at least 2 hours. Control plates equivalently supplemented with DMSO were prepared in parallel. L4 stage animals were shifted to drug or control DMSO-supplemented plates and imaged after 16-18 hours. Brightfield and fluorescence images of whole animals were collected on a Leica M165FC equipped with a 910 Leica K5 sCMOS camera and using LAS X software. Animals were immobilized for imaging using sodium azide and mounted on 2% agarose pads. All images were processed and analyzed using ImageJ/Fiji software. Images shown within the same figure panel were collected using the same exposure time and were then processed identically. To quantify accumulation of Ub(G76V)::GFP, the mean pixel intensity in a polygon selection of the anterior half of the intestine of each animal was measured. To quantify *rpt-3_p_::GFP* reporter expression, GFP::RPN-8 levels, and PBS-4::GFP levels, the mean pixel intensity was measured using a polygon selection around each animal.

### RNA Sequencing and Analysis

To prepare samples, 500 µL of synchronized L1s at a concentration of 80 worms/µL were placed in a 50 mL Falcon tube containing 24 mL NGM with *E. coli* HB101 (OD600 of 1.8-2.2) and 5.5 mL M9 for a total volume of 30 mL. Six conditions were tested: 1% DMSO control, 7.5 μM selectivin, 7.5 μM tioxazafen, 1 mM 1-ABT control, 1 mM 1-ABT + 7.5 μM selectivin, and 1 mM 1-ABT + 7.5 μM tioxazafen. Two tubes were prepared for each condition as technical replicates and three biological replicates were conducted on different days. Worms were pre-incubated for 4 hours with DMSO or 1 mM 1-ABT at 20°C on a nutator. Subsequently, DMSO or selectivin/tioxazafen was added and worms were incubated for an additional 18 hours under the same conditions. Following incubation, worms were pelleted by centrifugation at 2000 RPM for 1 minute (Eppendorf, P/N 5424), supernatant was removed, and worms were washed with sterile M9 10 times to remove bacteria and compounds. Worms were transferred to 1.5 mL microcentrifuge tubes and centrifuged at 3500 RPM for 3 minutes (Eppendorf, P/N 5424). All surfaces and equipment were decontaminated with RNase Away (ThermoFisher) before further processing. 1 mL of Trizol (Invitrogen, P/N 12183555) was added per 100 µL of worm pellet and then samples were vortexed for 30 seconds, flash-frozen in liquid nitrogen, and thawed at 37°C with vortexing and freeze-thawing repeated six times. RNA was extracted using the Trizol Plus RNA Purification Kit (Invitrogen, P/N 12183555) and treated with the TURBO DNase (Invitrogen, P/N AM2238) to remove genomic DNA contamination. RNA quality was assessed using a NanoDrop spectrophotometer, with acceptable 260/280 ratios exceeding 2.0. Samples were stored at -80°C until submission for sequencing at the Centre for Applied Genomics (TCAG) at The Hospital for Sick Children (SickKids, Toronto). Samples were tested with a Bioanalyzer for quality control. Stranded poly(A) mRNA libraries were prepared and sequenced with paired-end 150 bp reads using the Illumina NovaSeq X platform. The sequence was analyzed and relevant comparisons of sample reads to produce LogFC and FDR data was performed by TCAG. Briefly, the quality of the sequences in FASTQ format was assessed using the FastQC (v.0.11.5) tool. Adapter removal and trimming of read ends with Phred quality score less than 25 were carried out using Trim Galore (v.0.5.0) and Cutadapt (v.1.10). Trimmed reads less than 40 bp in length were excluded from downstream analysis. The reads were screened for the presence of rRNA sequences using FastQ-Screen (v.0.10.0). RSeQC package (v.2.6.2) was used to assess read distribution, positional read duplication, gene body coverage, junction saturation and to confirm the strandedness of the reads. The reads were aligned to the *Caenorhabditis elegans* reference genome (GCA_000002985.3) with Ensembl release 111 annotations using the STAR aligner (v.2.6.0.c). STAR alignments were processed to extract raw read counts for genes using htseq-count (v.0.6.1p2). Differential gene expression results were obtained with edgeR (v.3.28.1) using R (v.3.6.1). Minimal initial filtering of 50 read counts per gene in at least two samples was applied to the dataset. For the edgeR analyses, read counts were normalized with the scaling factors from the Trimmed means of M-values (TMM) method and glmLRT functionality was used for differential analysis. The resulting p-value results were adjusted using Benjamini and Hochberg’s method for controlling false-discovery rate (FDR). The log2 fold changes and adjusted p-values were used for the selection of differentially expressed genes.

### *C. elegans skn-1a(mg570)* and SKN-01[DDDD] Viability Assay

A bacterial suspension in liquid NGM was prepared by concentrating a saturated overnight HB101 culture (OD_600_: 1.8-2.2). 40 μL of this bacterial suspension, containing either 1.25 mM 1-ABT or DMSO vehicle, was added to each well of a 96-well flat-bottom culture plate. Approximately 20 synchronized L1 N2 *wild type* or GR2245 *skn-1a(mg570)* worms were then added to each well in 10 µl of M9 buffer, achieving a final concentration of 1mM 1-ABT. The samples were incubated in the 1-ABT/DMSO-containing medium for 24 hours. Thereafter, 0.3 µl of the serially diluted tioxazafen or bortezomib compounds were pinned into plates containing worms from a multiwell plate using a 96-well pinning tool (V&P Scientific Inc). Plates with worm samples was sealed with parafilm and placed in a box with several wet paper towels to prevent evaporation. The worms were incubated at 20°C with shaking at 200 rpm. After 6 days, the number of viable adults and larvae was counted manually using a dissection microscope. Dead worms were identified as those that did not move upon vigorous agitation of the plate and may be optically clearer with disrupted internal structures.

For the analysis of SKN-1A[DDDD], NGM plates containing the desired concentration of tioxazafen (Aaron Chemicals, #AR01QXD0) were prepared by directly applying tioxazafen solution to NGM plates seeded with OP50 and leaving the plates to dry for at least 2 hours. To measure survival, 20 L4-stage animals were transferred to fresh plates supplemented with 10 μM tioxazafen and incubated at 20°C. Survival of adult animals was scored after 48 hours.

### *In-vitro* Proteasome Activity Assay

80,000 synchronized L1s were incubated for 18 hours in 5 mL culture media in 15 mL-polypropylene tubes (FroggaBio Scientific Solutions, Ontario, Canada) containing *E. coli* HB101 bacteria (OD_600_ = 2.0-2.2) and liquid NGM, to which either 7.5 µM Tioxazafen in DMSO or DMSO (solvent control) was added with DMSO being a final concentration of 1% in all samples. Worms were lysed according to published work (Eroglu *et al*. 2024) with the following modifications. Worms were pelleted by centrifugation at 2,000 RPM for 1 minute (IEC Centra CL3R, Thermo Electron Corp., Massachusetts, U.S.A.) and washed several times with M9 buffer to remove bacteria. After the last wash, the buffer was aspirated to a minimum volume of 50-150 µL. The pellet was carefully transferred to a sterile Eppendorf tube to which was added 4 volumes of worm lysis buffer (50 mM HEPES, pH 7.5, 150 mM NaCl and 0.5% NP-40). The mixture was agitated vigorously for 7 minutes using a homogenizer (Bead Ruptor 24 Bead Mill Homogenizer, Omni International, Georgia, U.S.A.) followed by water sonication (5 minutes, 30 s ON/OFF cycles) (Symphony 97043-936 Ultrasonic Cleaner, VWR, Pennsylvania, U.S.A.). Worm debris was separated using centrifugation at 4°C for 10 minutes using a table-top centrifuge (Sorvall Legend Micro 21R, Thermo Scientific, Massachusetts, U.S.A.). Clear lysate was transferred carefully to fresh sterile Eppendorf tube placed on ice. If not used on the same day, the freshly-prepared worm lysate was immediately flash-frozen in liquid nitrogen and stored at -80°C and used within 4-5 days. Total protein concentration was quantified by the BCA protein quantification assay (Pierce BCA Protein Assay Kit, Thermo Scientific, Massachusetts, U.S.A.). Ten micrograms of total protein per sample was assayed using 96-well black, flat, clear-bottomed Nunc plates (Thermo Scientific, Massachusetts, U.S.A.) using Abcam proteasomal activity assay kit (Abcam ab107921) according to manufacturer’s instructions, with post-lysis additions of 1% DMSO or 7.5 µM Tioxazafen or 15 µM bortezomib. Fluorescence was measured every 10 minutes for 12 hours at a set temperature of 25°C using CLARIOstar plate-reader (BMG Labtech, Ortenberg, Germany). The experiment was repeated 3-7 times. Statistical analyses were performed on GraphPad Prism 10 (Boston, MA 02110, USA) using non-parametric Mann-Whitney U-test (p<0.05).

### Human Cell Culture and Stable Cell Line Generation

HEK293T (female, ATCC CRL-3216) and Lenti-X™ 293T (female, Clontech 632180) cells were cultured in Dulbecco’s Modified Eagle Medium (DMEM) (319-005 CL) supplemented with 10% fetal bovine serum (FBS) (Gibco, A5256701) and 1% penicillin-streptomycin (PS) (Multicell, 450-201-EL). Cell were maintained in a humidified incubator at 37°C and 5% CO2. Cell cultures were split for maintenance by washing with Dulbecco’s Phosphate Buffered Saline (D-PBS) (Multicell 311-425-CL) then treating with 0.25% Trypsin-EDTA solution (Multicell 325-045-EL) to dissociate the cells for resuspension in fresh media. Cells were routinely tested for mycoplasma contamination. Entry clones of human P450s CYP1A1, CYP1B1, and CYP3A4 from the human ORFeome collection (v8.1/9.1) were cloned to add a stop codon, then into the lentiviral Gateway-compatible destination vector pLX301 (Addgene, plasmid 25895) (Yang *et al*. 2011). Lentiviral particles containing the resulting plasmids was produced by transfecting Lenti-X™ 293T cells with the plasmids pLX301-[P450], psPAX2 (Addgene, plasmid 12260), and pVSV-G (Addgene, plasmid 8454) at a ratio of 5:3.5:1. Transfection was performed with polyethylenimine (PEI) "MAX" (Polysciences, cat #24765) transfection reagent at a ratio of 3µg PEI to 1µg DNA, suspended in Opti-MEM (Gibco 31985-070). The transfection mixture was vortexed and incubated at room temperature for 20 minutes before being added dropwise to cells. 16 hours post-transfection, the media was replaced with fresh DMEM. 48 hours post-transfection, the media was collected and filtered through a 0.45 µM filter. The lentivirus-containing supernatant was either aliquoted into cryotubes and frozen at -80°C for later use or was used immediately to transduce cells. To generate the HEK293T P450-expressing stable cell lines, cells were transduced with lentivirus containing the corresponding pLX301-[P450] vector at a high MOI in the presence of 8µg/mL polybrene (Sigma-Aldrich 107689-10G). 1.5µg/mL of puromycin (Gibco A11138-03) was added to cells 48 hours post-transduction to select for successfully transduced cells. Cells were selected for 48 hours or until an identically treated sample of un-transduced cells was completely dead.

### HEK293T Dose-Response Cell Viability Experiments

P450-expressing HEK293T stable cell lines were seeded in 96-well culture format at 3,500 cells per well. 24 hours later, they were treated with a titration of tioxazafen or benzo[a]pyrene, normalizing each well to the same concentration of DMSO, not exceeding a final volume of 1% DMSO. Cells were cultured for another 48 hours, then CellTiter-Glo® Luminescent Cell Viability Assay reagent (Promega, G7573) was added onto cells to lyse them and detect the number of viable cells, per the manufacturer’s instructions. Luminescence intensities were measured using a multimode microplate reader (Biotek). Luminescence intensity was normalized to the DMSO-only treatment, and the resulting cell viability values were plotted with GraphPad Prism 10 with a variable-slope, four-parameter, non-linear regression curve fit.

### HEK293T RNA-seq

P450-expressing HEK293T stable cell lines were seeded in 24-well culture format, 60,000 cells per well. 24 hours later, they were treated with tioxazafen or DMSO. 24 hours post-treatment, they were trypsinized with TrypLE™ Express Enzyme (Gibco, 12604021) to dissociate them from the plate. 100,000 to 200,000 cells were centrifuged to pellet them, then washed with phosphate-buffered saline, and finally centrifuged to pellet again. Cells were resuspended in 50µL of Zymo Research Corporation DNA/RNA Shield™ (R1100-50). RNA-seq was performed by Plasmidsaurus using Illumina Sequencing Technology with custom analysis and annotation.

### HEK293T Sample Preparation for LC-MS

P450-expressing HEK293T stable cell lines were seeded in 6-well culture format, 350,000 cells per well. 48 hours later, they were treated with tioxazafen or DMSO and media was replaced with Opti-MEM supplemented with 10% FBS. 4 hours later, cells were collected by a cell scraper, washed with PBS, and pelleted. Samples were normalized by cell pellet weight, approximately 1,000,000 cells, and frozen at -80°C before processing.

