## Supplemental Display Items for "The Nematicide Tioxazafen Disrupts Proteasome Function via Cytochrome P450 Bioactivation"

### Supplemental Table 1

| Dataset | Source | Term | p-value | fold change | Fraction of input gene list annotated to a bioprocess cluster | Cluster frequency | Background frequency |
|---|---|---|---|---|---|---|---|
| 33 selective enhancers of bioactivated tioxazafen | this work | Protein turnover | 4.2E-09 | 9.79 | 11 / 31, 35.5% | 4 / 11, 36.4% | 65 / 1750, 3.7% |
| | | Mitosis | 3.6E-05 | 3.28 | 11 / 31, 35.5% | 2 / 11, 18.2% | 97 / 1750, 5.5% |
| | | Cell polarity | 3.6E-05 | 2.49 | 11 / 31, 35.5% | 2 / 11, 18.2% | 128 / 1750, 7.3% |
| | | Glycosylation & Protein folding | 2.1E-04 | 1.61 | 11 / 31, 35.5% | 2 / 11, 18.2% | 198 / 1750, 11.3% |
| 163 selective enhancers of bioactivated selectivin | this work | Glycosylation & Protein folding | 1.8E-42 | 3.79 | 63 / 151, 41.7% | 27 / 63, 42.9% | 198 / 1750, 11.3% |
| | | Vesicle traffic | 2.0E-20 | 3.06 | 63 / 151, 41.7% | 14 / 63, 22.2% | 127 / 1750, 7.3% |
| | | DNA replication & repair | 1.4E-03 | 0.33 | 63 / 151, 41.7% | 3 / 63, 4.8% | 251 / 1750, 14.3% |
| | | Mitosis | 3.0E-03 | 0.86 | 63 / 151, 41.7% | 3 / 63, 4.8% | 97 / 1750, 5.5% |
| | | Protein turnover | 7.1E-03 | 0.85 | 63 / 151, 41.7% | 2 / 63, 3.2% | 65 / 1750, 3.7% |
| | | Transcription | 7.1E-03 | 0.54 | 63 / 151, 41.7% | 3 / 63, 4.8% | 154 / 1750, 8.8% |
| | | Cell polarity | 1.7E-02 | 0.43 | 63 / 151, 41.7% | 2 / 63, 3.2% | 128 / 1750, 7.3% |
| | | Cytokinesis | 6.9E-02 | 1.85 | 63 / 151, 41.7% | 1 / 63, 1.6% | 15 / 1750, 0.9% |
| 47 enhancers of 1.3 mM bortezomib (SGA score of <-3.0) | Berry et al., 2011 | Protein turnover | 7.6E-14 | 11.54 | 14 / 42, 33.3% | 6 / 14, 42.9% | 65 / 1750, 3.7% |
| | | Transcription | 5.8E-04 | 1.62 | 14 / 42, 33.3% | 2 / 14, 14.3% | 154 / 1750, 8.8% |
| | | Cell polarity | 8.0E-03 | 0.98 | 14 / 42, 33.3% | 1 / 14, 7.1% | 128 / 1750, 7.3% |
| | | Ribosome biogenesis | 1.6E-02 | 1.34 | 14 / 42, 33.3% | 1 / 14, 7.1% | 93 / 1750, 5.3% |
| 64 enhancers of 2.5 mM DTT (SGA score of <-3.0) | Berry et al., 2011 | Glycosylation & Protein folding | 6.25E-17 | 5.05051 | 14 / 60, 23.3% | 8 / 14, 57.1% | 198 / 1750, 11.3% |
| | | Vesicle traffic | 1.28E-08 | 3.93701 | 14 / 60, 23.3% | 4 / 14, 28.6% | 127 / 1750, 7.3% |
| | | DNA replication & repair | 0.001206 | 0.99602 | 14 / 60, 23.3% | 2 / 14, 14.3% | 251 / 1750, 14.3% |

**Supplemental Table 1. Gene Categories Enriched within the Yeast Genetic Interaction Network Among the Mutant Genes that Enhance the Phenotype of Bioactivated Tioxazafen and Controls.** The significant ($p<0.05$) enrichment scores from Spatial Analysis of Functional Enrichment (SAFE) analyses (Baryshnikova, 2016) obtained from the online Cell Map tool (Usaj *et al.*, 2017).

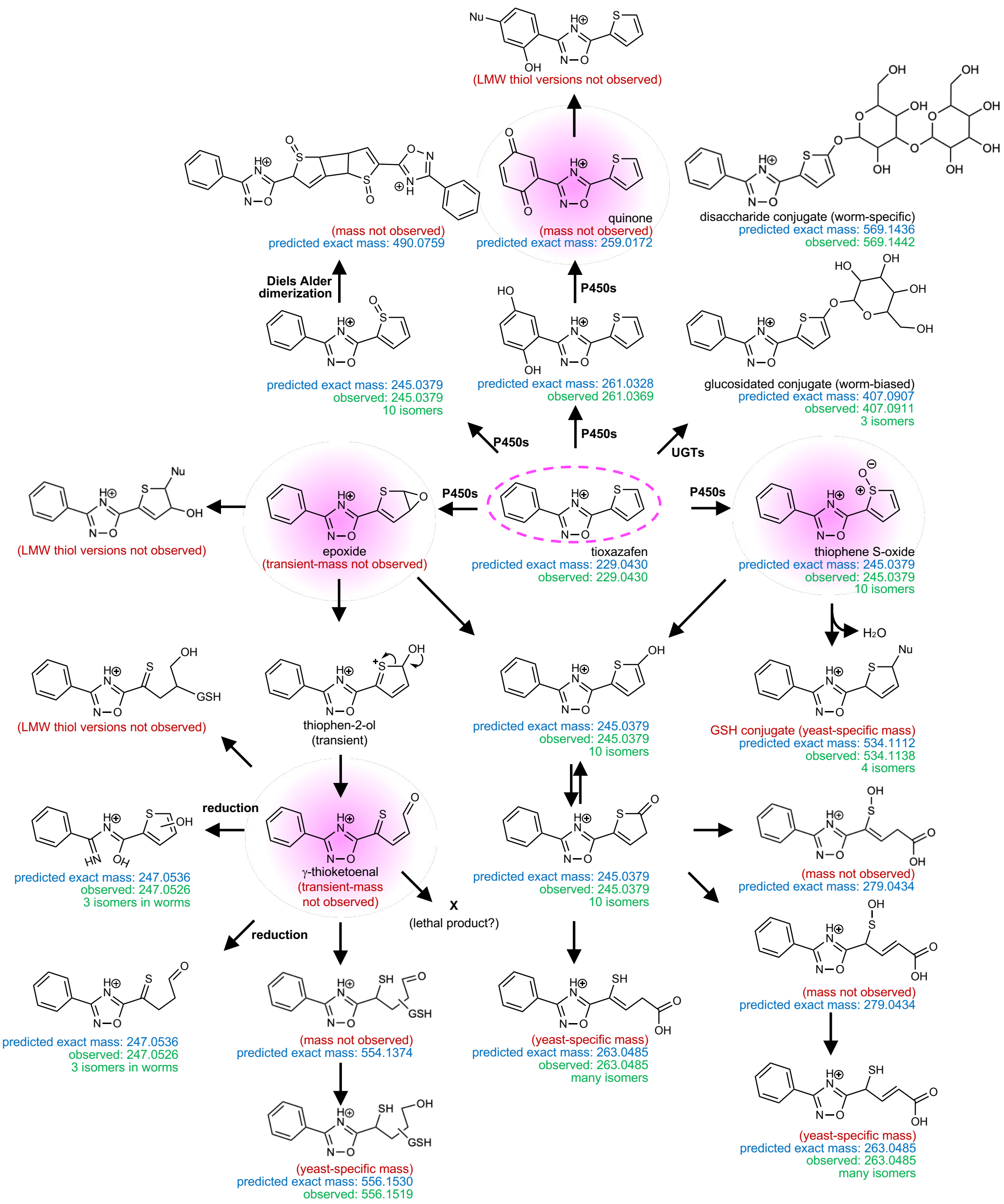


**Supplemental Figure 1. Theoretical Routes for Tioxazafen Bioactivation Along with Exact [H$^+$] Masses.** The reaction schemes are largely based on Gramec *et al*., 2014 and Podgorski *et al*., 2023. Shown are the predicted structures (and their masses) based on the observed exact masses. The four predicted reactive structures are highlighted with a pink background. The epoxide/g-thioketoenal pathway that leads to the 247.0526 m/z observed masses track with lethality in both worms and yeast, consistent with the idea that this pathway leads to a product that disrupts proteasome function. All the other routes that generate reactive theoretical metabolites lack evidence of the route's existence in worms, although the reactive products could be rapidly consumed by targets and are not discounted. Note: i) we lack confidence in the existence of the di-hydroxylated metabolite (exact mass 261.0369 highlighted in yellow) because of the departure from the expected mass, ii) the depicted disaccharide conjugate is unprecedented, iii) the conjugation site for the glucosidated and disaccharide conjugates are speculative.

### Supplemental Figure 2

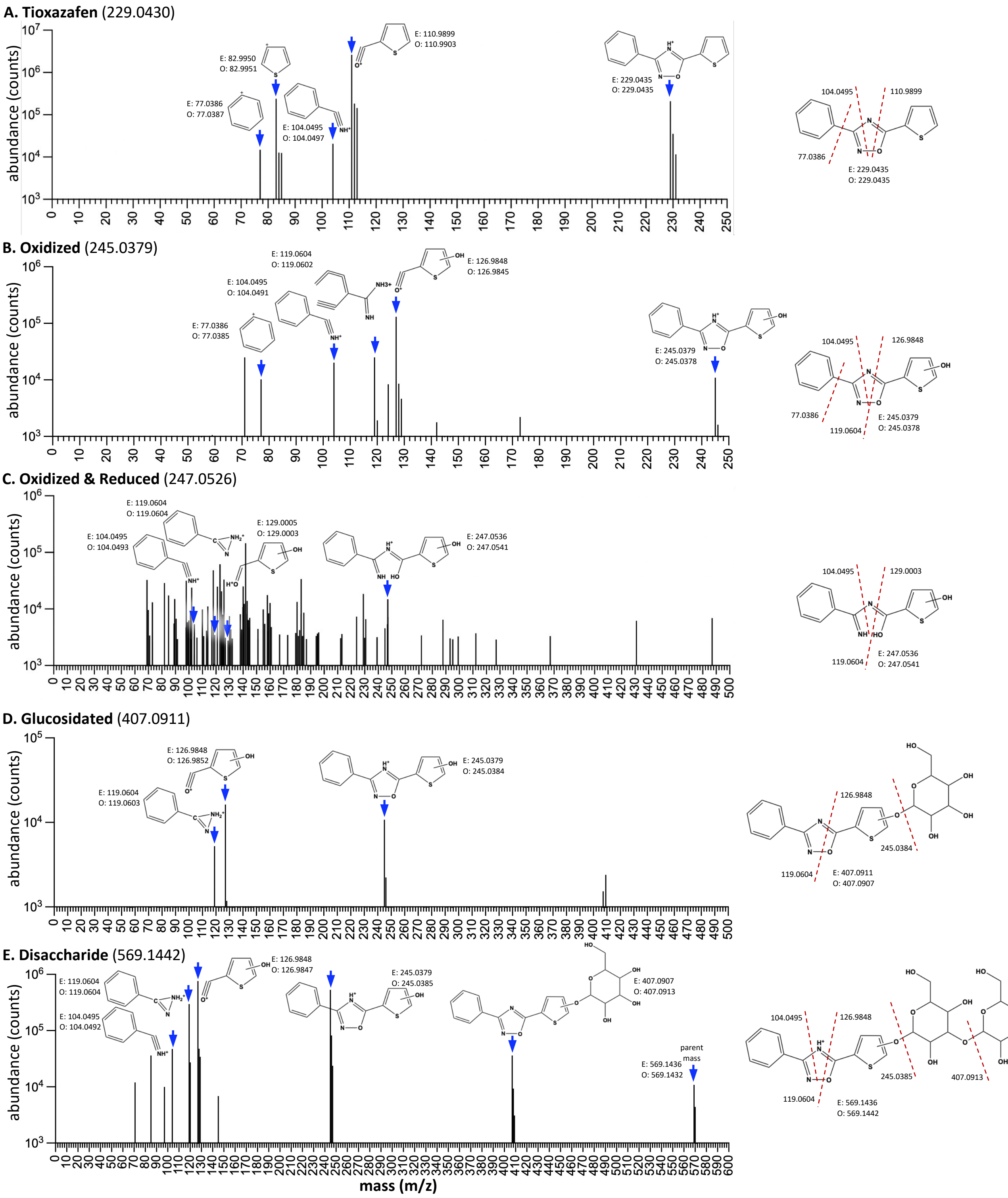


**Supplemental Figure 2. Tandem Mass-Spec Analyses of *C. elegans*-Generated Tioxazafen Metabolites.** All masses fragmented come from *C. elegans* lysates incubated with Tioxazafen (100 μM) for 24 hours. All structures shown are predictions based on exact masses; there may be alternative structures based on the exact mass (e,g, see the two alternative structures provided for 119.0604in B and C). How the metabolite or parent molecule likely fragments based on the exact masses is shown on the right with a dotted red line. The exact position of the OH group and sugars is not known except that the fragmentation patterns indicate that it is likely on the thiophene ring. E, expected exact mass based on structure; O, observed exact mass. Note that three different collision energies were used (see methods) and fragment data from each collision energy is combined in each plot. The source exact mass from the original run (done many days prior to the tandem mass spectrometry experiment) is shown in brackets in the subheading of each panel. See the DataFile for additional details.

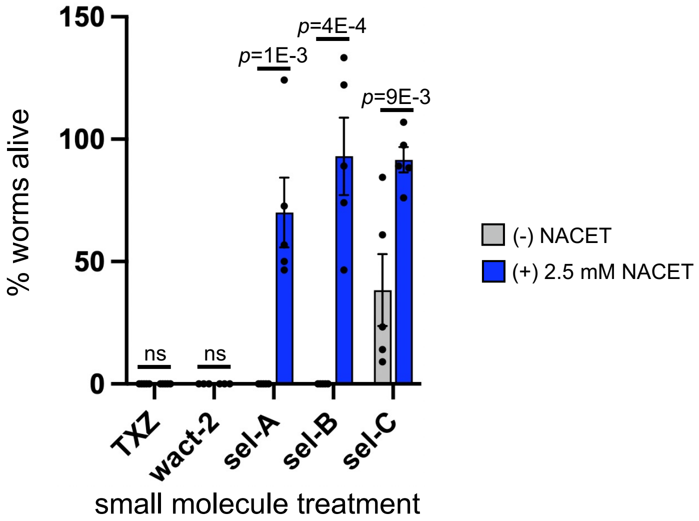


**Supplemental Figure 3. NACET Suppresses the Lethal Effects of Selectivin and its Analogs, but Not Tioxazafen or wact-2 .** Wild type (N2) first larval stage (L1) animals were treated with NACET or negative control for 4 hours before incubation with drugs. The animals were then treated for 24 hours with 100 micromolar of the indicated drug. The percent of worms alive relative to the DMSO control was calculated based on the number of animals moving in the experimental samples relative to the DMSO controls. At least three independent trials were performed. Standard error of the mean is shown. A two-tailed Student's T-test was used to analyze significant differences. TXZ, tioxazafen; sel, selectivin; ns, not significant.

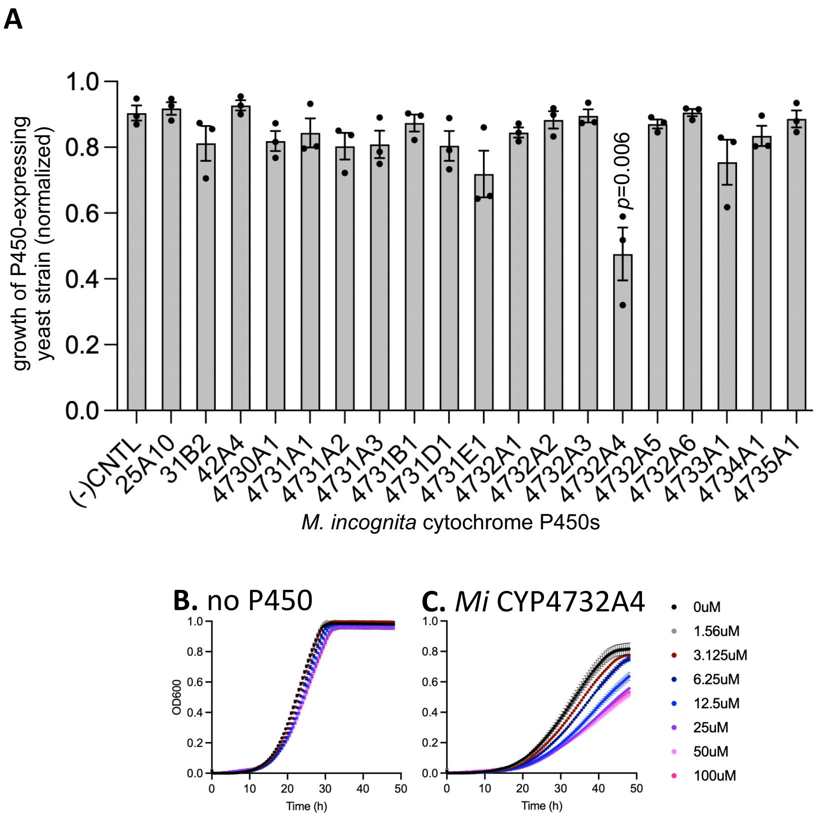


**Supplemental Figure 4. A.** A PIXY survey of the ability of 19 *Meloidogyne incognita* P450s, when heterologously expressed in yeast, to bioactivate tioxazafen (50 μM) into a lethal product. Three independent repeats were done with two technical replicates each (N=3, n=2). **B-C.** Dose-response analyses of tioxazafen in yeast strains expressing the indicated P450s.

### Supplemental Figure 5

| gene name | GFP reporter expression in response to tioxazafen |
|---|---|
| gst-4 | 0.8 |
| sod-3 | 0.5 |
| irg-1 | 1.3 |
| nlp-29 | 1.0 |
| cnc-2 | 0.8 |
| gpdh-1 | 2.5 |
| hsp-4 | 2.3 |
| hsp-6 | 1.8 |
| hsp-16.41 | 2.5 |
| hsp-70 | 2.8 |
| hsp-90 | 2.8 |
| rpt-3 | 3.0 |

| gene | tioxazafen RNAseq | | | | selectivin RNAseq | | | |
|---|---|---|---|---|---|---|---|---|
| | parent + met | | parent | | parent + met | | parent | |
| name | logFC | FDR | logFC | FDR | logFC | FDR | logFC | FDR |
| gst-4 | 2.9 | 1.E-15 | 1.3 | 3.E-04 | 2.8 | 7.E-14 | 2.2 | 3.E-09 |
| sod-3 | -0.3 | 4.E-01 | 0.6 | 4.E-02 | -0.1 | 9.E-01 | 0.9 | 2.E-03 |
| irg-1 | 2.5 | 3.E-09 | 0.5 | 4.E-01 | 1.7 | 2.E-04 | 0.2 | 8.E-01 |
| nlp-29 | 0.8 | 3.E-02 | -0.5 | 3.E-01 | 0.5 | 3.E-01 | -0.5 | 2.E-01 |
| cnc-2 | -2.9 | 1.E-07 | -1.3 | 5.E-03 | 1.8 | 1.E-04 | 0.3 | 6.E-01 |
| gpdh-1 | -0.6 | 1.E-01 | -0.3 | 6.E-01 | -0.9 | 4.E-02 | -0.4 | 4.E-01 |
| hsp-4 | 2.2 | 1.E-12 | -0.3 | 4.E-01 | -0.2 | 7.E-01 | -0.4 | 3.E-01 |
| hsp-6 | 1.3 | 1.E-12 | 1.0 | 2.E-07 | 0.7 | 6.E-04 | 0.7 | 5.E-04 |
| hsp-16.41 | 3.4 | 2.E-12 | 1.9 | 9.E-05 | 0.9 | 1.E-01 | 1.8 | 4.E-04 |
| hsp-70 | 2.5 | 6.E-08 | 2.0 | 2.E-05 | 0.5 | 4.E-01 | 2.1 | 2.E-05 |
| hsp-90 | 1.2 | 9.E-10 | 0.6 | 6.E-03 | 0.2 | 4.E-01 | 0.6 | 3.E-03 |
| rpt-3 | 1.5 | 6.E-29 | 0.0 | 9.E-01 | 0.0 | 9.E-01 | 0.1 | 8.E-01 |

0.0 0.5 1.0 1.5 2.0 2.5 3.0

GFP Reporter Expression (AU)

-3.0 -2.5 -2.0 -1.5 -1.0 -0.5 0.0 0.5 1.0 1.5 2.0 2.5 3.0

RNAseq fold regulated ($log_2$)

$\leq$1E-7 1.E-07 1.E-06 1.E-05 1.E-04 1.E-03 $\leq$1E-3

RNAsqq false discovery rate (FDR)

**Supplemental Figure 5. A Comparison of GFP reporter expression and RNAseq Data.** The GFP reporter column is the average score of the GFP expression from the relevant GFP reporter strain (N=1, n=2). The RNAseq columns show the log fold-change (LogFC) and FDR for the indicated categories (N=3, n=3 for each replicate). We can infer responses due to bioactivated tioxazafen (and selectivin) by comparing the transcriptional response elicited in the ‘parent + metabolites’ (met) columns (derived from the drug vs solvent control samples) to the just the ‘parent’ columns (derived from the drug +1-ABT vs 1-ABT alone samples). See the main text and the RNAseq methods for more details.

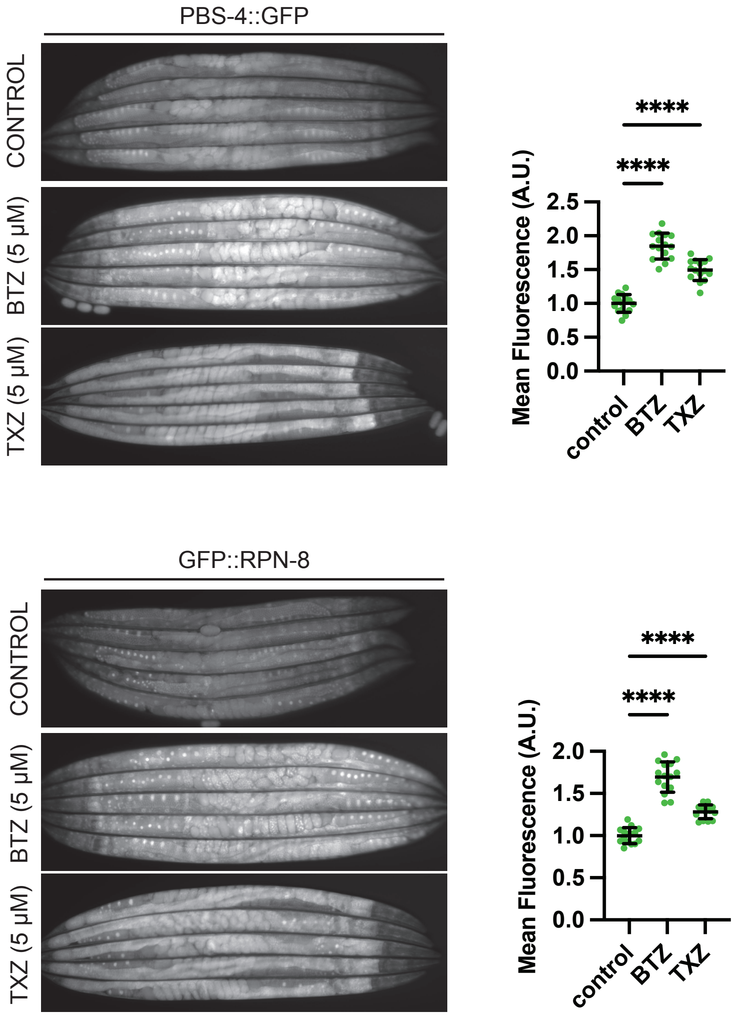


**Supplemental Figure 6. Tioxazafen Upregulates Expression of Proteosome Translational Reporters.** L4 animals were shifted to plates supplemented with either 5μM bortezomib, 5 μM tioxazafen, or DMSO control and incubated at 20°C for 18 hours before imaging. Quantification of n=15 animals per condition is shown. Error bars show mean ±SD. Significance was determined by ordinary one-way ANOVA with Dunnett's multiple comparisons test (**** p<0.0001).

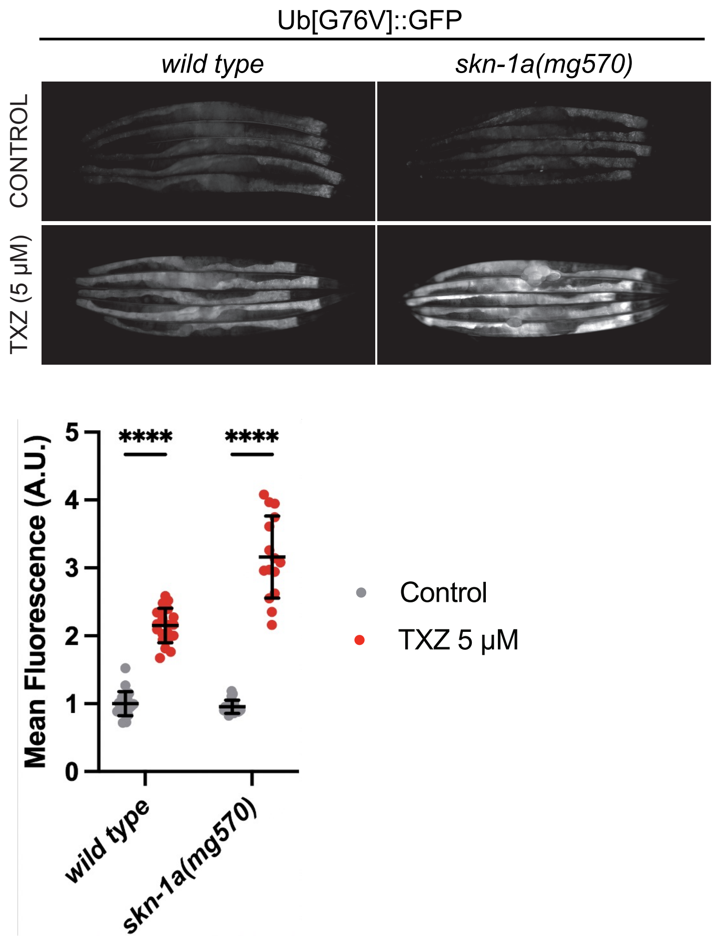


**Supplemental Figure 7. The Increase in UbV-GFP that is Induced by Tioxazafen is Compensated in Part by the Activity of SKN-1A.** L4 animals were shifted to plates supplemented with 5 µM tioxazafen or vehicle control and incubated at 20°C for 16 hours before imaging. Quantification of n≥15 animals per condition is shown. Error bars show mean ±SD. Significance was determined by ordinary two-way ANOVA with uncorrected Fisher's LSD test (**** $p<0.0001$).

**A**

| gene name | human ortholog | tioxazafen parent + met logFC | tioxazafen parent + met FDR | tioxazafen parent logFC | tioxazafen parent FDR | selectivin parent + met logFC | selectivin parent + met FDR | selectivin parent logFC | selectivin parent FDR |
|---|---|---|---|---|---|---|---|---|---|
| **Cyclin A Complex (prep for mitosis including kinetochore construction)** | | | | | | | | | |
| cya-1 | Cyclin A | -1.5 | 1.E-18 | -0.7 | 9.E-05 | -0.3 | 1.E-01 | -0.5 | 1.E-02 |
| cdk-1 | CDK1 | -3.4 | 4.E-61 | -1.5 | 1.E-13 | -0.6 | 5.E-03 | -0.7 | 6.E-04 |
| efl-1 | E2F | -0.7 | 3.E-05 | -0.5 | 5.E-03 | -0.3 | 2.E-01 | -0.4 | 4.E-02 |
| efl-2 | E2F | 0.5 | 1.E-04 | 0.3 | 5.E-02 | 0.3 | 1.E-01 | 0.1 | 5.E-01 |
| efl-3 | E2F | -3.4 | 2.E-77 | -1.1 | 1.E-10 | -1.0 | 3.E-08 | -0.8 | 2.E-05 |
| dpl-1 | TFDP1/2/3 | -1.1 | 1.E-11 | -0.2 | 4.E-01 | 0.1 | 8.E-01 | -0.1 | 6.E-01 |
| air-1 | Aurora A | -2.8 | 7.E-59 | -0.9 | 1.E-07 | -0.6 | 9.E-04 | -0.6 | 1.E-03 |
| air-2 | Aurora A | -3.1 | 1.E-35 | -1.5 | 2.E-10 | -0.7 | 5.E-03 | -0.8 | 2.E-03 |
| plk-1 | PLK1 | -2.5 | 1.E-44 | -0.8 | 2.E-06 | -0.4 | 3.E-02 | -0.5 | 1.E-02 |
| plk-2 | PLK1 | -3.5 | 7.E-12 | -2.4 | 3.E-06 | 0.0 | 1.E+00 | -1.0 | 9.E-02 |
| plk-3 | PLK1 | 0.0 | 9.E-01 | -0.5 | 9.E-02 | -0.3 | 4.E-01 | -0.7 | 2.E-02 |
| **Cyclin B Complex (activates mitotic entry)** | | | | | | | | | |
| cyb-1 | Cyclin B | -2.6 | 2.E-44 | -1.4 | 2.E-15 | -0.6 | 2.E-03 | -1.0 | 6.E-09 |
| cyb-3 | Cyclin B | -3.4 | 9.E-65 | -1.2 | 1.E-09 | -0.5 | 2.E-02 | -0.6 | 2.E-03 |
| cdc-25.2 | CDC25C | -3.7 | 5.E-39 | -1.6 | 3.E-09 | -1.1 | 2.E-04 | -1.1 | 1.E-04 |
| **Cyclin D Complex (promotes G1-S phase)** | | | | | | | | | |
| cyd-1 | Cyclin D | -2.9 | 4.E-25 | -1.7 | 7.E-10 | -0.5 | 1.E-01 | -1.1 | 1.E-04 |
| cdk-4 | CDK4 | -1.1 | 4.E-05 | 0.0 | 1.E+00 | -0.1 | 9.E-01 | 0.1 | 8.E-01 |
| dpl-1 | TFDP1/2/3 | -1.1 | 1.E-11 | -0.2 | 4.E-01 | 0.1 | 8.E-01 | -0.1 | 6.E-01 |
| **Cyclin E Complex (promotes G1-S phase)** | | | | | | | | | |
| cye-1 | Cyclin E | -2.3 | 6.E-34 | -1.3 | 2.E-12 | -0.3 | 2.E-01 | -1.0 | 2.E-07 |
| cdk-2 | CDK2 | -0.6 | 1.E-03 | -0.4 | 5.E-02 | -0.1 | 7.E-01 | -0.1 | 8.E-01 |
| cdc-25.2 | CDC25A | -3.7 | 5.E-39 | -1.6 | 3.E-09 | -1.1 | 2.E-04 | -1.1 | 1.E-04 |
| aspm-1 | ASPM | -2.8 | 4.E-36 | -0.7 | 1.E-03 | -0.3 | 2.E-01 | -0.4 | 1.E-01 |
| skpt-1 | Skp2 | -2.6 | 1.E-20 | -1.5 | 1.E-08 | -0.2 | 7.E-01 | -0.8 | 8.E-03 |
| sel-10 | FBXW7 | -0.5 | 5.E-04 | -0.4 | 2.E-03 | -0.4 | 2.E-02 | -0.3 | 9.E-02 |
| bmk-1 | KIF11/p27 | -2.7 | 2.E-44 | -0.9 | 1.E-06 | -0.4 | 1.E-01 | -0.6 | 7.E-03 |
| lin-35 | Rb | -0.6 | 1.E-03 | -0.3 | 2.E-01 | -0.3 | 3.E-01 | -0.1 | 8.E-01 |
| dpl-1 | TFDP1/2/3 | -1.1 | 1.E-11 | -0.2 | 4.E-01 | 0.1 | 8.E-01 | -0.1 | 6.E-01 |

**B**

| gene name | human ortholog | tioxazafen parent + met logFC | tioxazafen parent + met FDR | tioxazafen parent logFC | tioxazafen parent FDR | selectivin parent + met logFC | selectivin parent + met FDR | selectivin parent logFC | selectivin parent FDR |
|---|---|---|---|---|---|---|---|---|---|
| **Cytokinesis-Cytokinetic Furrow** | | | | | | | | | |
| zen-4 | KIF23 | -2.5 | 4.E-51 | -0.8 | 3.E-06 | -0.4 | 6.E-02 | -0.4 | 2.E-02 |
| cyk-4 | RACGAP1 | -2.4 | 1.E-34 | -0.7 | 8.E-04 | -0.3 | 3.E-01 | -0.5 | 2.E-02 |
| unc-61 | Septin | -2.9 | 4.E-39 | -0.7 | 2.E-03 | -0.4 | 1.E-01 | -0.4 | 1.E-01 |
| unc-59 | Septin | -2.3 | 1.E-37 | -0.7 | 1.E-04 | -0.4 | 6.E-02 | -0.3 | 1.E-01 |
| ani-1 | Anillin | -1.8 | 5.E-16 | 0.0 | 1.E+00 | 0.0 | 1.E+00 | -0.1 | 8.E-01 |
| ani-2 | Anillin | -0.1 | 8.E-01 | 0.0 | 9.E-01 | -0.2 | 7.E-01 | -0.4 | 2.E-01 |
| ani-3 | Anillin | -0.8 | 8.E-04 | -0.2 | 5.E-01 | -0.3 | 3.E-01 | -0.2 | 5.E-01 |
| ect-2 | ECT2 | -2.3 | 3.E-31 | -0.5 | 2.E-02 | -0.2 | 4.E-01 | -0.4 | 1.E-01 |
| ocrl-1 | OCRL | -0.4 | 2.E-03 | -0.2 | 3.E-01 | -0.2 | 4.E-01 | -0.2 | 3.E-01 |
| rom-1 | RHBDL1 | 0.8 | 3.E-04 | -0.4 | 6.E-02 | -0.2 | 6.E-01 | -0.3 | 2.E-01 |
| cyk-1 | DIAPH2 | -1.2 | 2.E-11 | -1.0 | 1.E-07 | -0.7 | 7.E-04 | -1.0 | 1.E-07 |
| let-502 | ROCK1/2 | -1.2 | 2.E-10 | -0.6 | 3.E-03 | -0.2 | 3.E-01 | -0.5 | 3.E-02 |
| rhgf-1 | ARHGEF | -1.1 | 1.E-08 | -0.5 | 1.E-02 | -0.6 | 2.E-03 | -0.6 | 3.E-03 |
| pfn-3 | n/a | -1.3 | 4.E-10 | -0.4 | 7.E-02 | -0.5 | 3.E-02 | -0.2 | 4.E-01 |
| pfn-2 | n/a | 0.4 | 2.E-02 | -0.6 | 1.E-03 | -0.6 | 1.E-03 | -0.4 | 4.E-02 |

fold regulated ($\log_2$): ≤-2 | -2.0 | -1.5 | -1.0 | -0.5 | 0.0 | ≥0

false discovery rate (FDR): ≤1E-7 | 1.E-07 | 1.E-06 | 1.E-05 | 1.E-04 | 1.E-03 | ≤1E-3

**C**

| gene name | human ortholog | tioxazafen parent + met logFC | tioxazafen parent + met FDR | tioxazafen parent logFC | tioxazafen parent FDR | selectivin parent + met logFC | selectivin parent + met FDR | selectivin parent logFC | selectivin parent FDR |
|---|---|---|---|---|---|---|---|---|---|
| **Kinetochore Machinery** | | | | | | | | | |
| hcp-3 | CENP-A | -1.7 | 2.E-31 | -0.8 | 4.E-09 | -0.3 | 1.E-01 | -0.5 | 5.E-04 |
| knl-2 | n/a | -2.1 | 9.E-23 | -0.9 | 3.E-05 | -0.3 | 3.E-01 | -0.6 | 6.E-03 |
| hcp-4 | CENP-C | -2.0 | 3.E-28 | -0.9 | 4.E-06 | -0.3 | 2.E-01 | -0.6 | 3.E-03 |
| hcp-6 | NCAPD3 | -2.9 | 2.E-34 | -0.9 | 3.E-04 | -0.4 | 2.E-01 | -0.7 | 1.E-02 |
| san-1 | MAD3 | -2.0 | 2.E-12 | -0.8 | 5.E-03 | -0.1 | 9.E-01 | -0.3 | 3.E-01 |
| mel-28 | n/a | -0.8 | 8.E-06 | -0.4 | 4.E-02 | 0.0 | 1.E+00 | -0.4 | 7.E-02 |
| lis-1 | n/a | 0.1 | 7.E-01 | -0.1 | 4.E-01 | 0.0 | 9.E-01 | -0.2 | 3.E-01 |
| klp-7 | MCAK | -0.4 | 1.E-02 | -0.4 | 3.E-02 | 0.0 | 1.E+00 | -0.4 | 3.E-02 |
| rod-1 | KNTC1 | -1.9 | 6.E-16 | -0.5 | 7.E-02 | -0.2 | 6.E-01 | -0.3 | 3.E-01 |
| zwl-1 | ZWILCH | -1.7 | 2.E-14 | -0.8 | 3.E-04 | 0.0 | 9.E-01 | -0.5 | 6.E-02 |
| czw-1 | ZW10 | -0.7 | 5.E-05 | -0.4 | 2.E-02 | -0.1 | 7.E-01 | -0.2 | 3.E-01 |
| bub-1 | BUB1 | -2.8 | 2.E-38 | -0.9 | 2.E-05 | -0.4 | 1.E-01 | -0.6 | 2.E-02 |
| bub-3 | BUB3 | -1.2 | 3.E-11 | -0.5 | 2.E-02 | -0.1 | 6.E-01 | -0.3 | 1.E-01 |
| plk-1 | PLK1 | -2.5 | 1.E-44 | -0.8 | 2.E-06 | -0.4 | 3.E-02 | -0.5 | 1.E-02 |
| hcp-1 | CENP-F | -3.0 | 3.E-33 | -0.9 | 3.E-04 | -0.3 | 4.E-01 | -0.7 | 1.E-02 |
| hcp-2 | CENP-F | -2.6 | 1.E-37 | -1.0 | 4.E-07 | -0.3 | 2.E-01 | -0.6 | 3.E-03 |
| cls-2 | CLASP | 0.4 | 1.E-02 | -0.1 | 8.E-01 | 0.3 | 1.E-01 | 0.0 | 9.E-01 |
| mdf-1 | MAD1 | -0.1 | 5.E-01 | 0.0 | 8.E-01 | 0.0 | 9.E-01 | 0.1 | 6.E-01 |
| mdf-2 | MAD2 | -2.1 | 6.E-24 | -0.5 | 1.E-02 | -0.2 | 6.E-01 | -0.2 | 3.E-01 |
| knl-1 | KNL1 | -2.6 | 3.E-26 | -1.1 | 5.E-06 | -0.3 | 3.E-01 | -0.7 | 6.E-03 |
| kbp-5 | n/a | -1.9 | 3.E-14 | -0.7 | 6.E-03 | -0.2 | 7.E-01 | -0.3 | 4.E-01 |
| kbp-3 | SPBC25 | -1.6 | 8.E-10 | -1.1 | 2.E-05 | -0.1 | 8.E-01 | -0.6 | 3.E-02 |
| kbp-4 | SPBC24 | -3.0 | 6.E-21 | -1.3 | 1.E-05 | -0.2 | 6.E-01 | -0.4 | 2.E-01 |
| him-10 | NUF2 | -2.3 | 6.E-28 | -1.1 | 3.E-07 | -0.2 | 6.E-01 | -0.5 | 2.E-02 |
| ndc-80 | NDC80 | -1.6 | 9.E-17 | -0.6 | 3.E-03 | -0.1 | 8.E-01 | -0.2 | 3.E-01 |
| mis-12 | MIS12 | -2.8 | 7.E-39 | -1.1 | 8.E-08 | -0.5 | 3.E-02 | -0.6 | 1.E-02 |
| kbp-1 | n/a | -3.0 | 9.E-15 | -1.5 | 3.E-05 | -0.2 | 7.E-01 | -0.6 | 1.E-01 |
| kbp-2 | n/a | -2.5 | 4.E-20 | -1.0 | 2.E-04 | -0.3 | 4.E-01 | -0.3 | 4.E-01 |
| knl-3 | n/a | -2.2 | 6.E-28 | -0.9 | 2.E-06 | -0.2 | 5.E-01 | -0.5 | 2.E-02 |
| ska-1 | SKA1 | -0.2 | 2.E-01 | -0.2 | 2.E-01 | 0.1 | 6.E-01 | -0.1 | 8.E-01 |
| ska-3 | n/a | -1.6 | 2.E-15 | -1.0 | 1.E-06 | -0.3 | 3.E-01 | -0.3 | 2.E-01 |
| klp-19 | KIF4A | -2.4 | 1.E-31 | -0.9 | 3.E-05 | -0.2 | 4.E-01 | -0.6 | 8.E-03 |
| spd-1 | PRC1 | -1.7 | 6.E-22 | -0.7 | 5.E-04 | -0.1 | 8.E-01 | -0.3 | 2.E-01 |
| tba-1 | α-tubulin | -1.3 | 1.E-14 | -0.5 | 1.E-02 | -0.1 | 7.E-01 | -0.3 | 2.E-01 |
| tba-8 | α-tubulin | -4.7 | 2.E-23 | -1.6 | 2.E-04 | -0.3 | 6.E-01 | -0.7 | 2.E-01 |
| tbb-1 | β-tubulin | -0.6 | 1.E-03 | -0.1 | 8.E-01 | 0.0 | 1.E+00 | 0.1 | 8.E-01 |
| tbb-2 | β-tubulin | -0.9 | 1.E-06 | -0.5 | 1.E-02 | -0.2 | 6.E-01 | -0.4 | 7.E-02 |

**D**

| gene name | human ortholog | tioxazafen parent + met logFC | tioxazafen parent + met FDR | tioxazafen parent logFC | tioxazafen parent FDR | selectivin parent + met logFC | selectivin parent + met FDR | selectivin parent logFC | selectivin parent FDR |
|---|---|---|---|---|---|---|---|---|---|
| **spindle duplication** | | | | | | | | | |
| spd-2 | n/a | -0.3 | 1.E-01 | -0.4 | 2.E-02 | 0.3 | 9.E-02 | -0.2 | 2.E-01 |
| zyg-1 | n/a | -0.7 | 2.E-04 | -0.2 | 4.E-01 | 0.1 | 7.E-01 | -0.3 | 2.E-01 |
| sas-6 | n/a | -1.9 | 9.E-29 | -0.7 | 1.E-05 | -0.2 | 3.E-01 | -0.4 | 3.E-02 |
| sas-5 | n/a | -1.4 | 5.E-15 | -0.8 | 2.E-06 | -0.1 | 7.E-01 | -0.5 | 9.E-03 |
| sas-4 | n/a | -1.3 | 2.E-19 | -0.6 | 1.E-04 | -0.3 | 9.E-02 | -0.5 | 4.E-03 |
| sas-7 | n/a | 0.5 | 1.E-03 | -0.3 | 7.E-02 | 0.1 | 7.E-01 | -0.3 | 7.E-02 |
| air-1 | Aurora A | -2.8 | 7.E-59 | -0.9 | 1.E-07 | -0.6 | 9.E-04 | -0.6 | 1.E-03 |
| air-2 | Aurora A | -3.1 | 1.E-35 | -1.5 | 2.E-10 | -0.7 | 5.E-03 | -0.8 | 2.E-03 |
| spd-5 | n/a | -1.8 | 1.E-25 | -0.4 | 3.E-02 | -0.2 | 3.E-01 | -0.2 | 2.E-01 |
| tbg-1 | γ-tubulin | -0.5 | 2.E-04 | 0.0 | 9.E-01 | 0.0 | 1.E+00 | 0.1 | 7.E-01 |
| tac-1 | n/a | -0.3 | 8.E-02 | -0.6 | 5.E-04 | -0.1 | 8.E-01 | -0.2 | 2.E-01 |
| zyg-9 | CKAP5 | -0.5 | 7.E-03 | -0.4 | 3.E-02 | 0.1 | 8.E-01 | -0.4 | 8.E-02 |
| klp-7 | MCAK | -0.4 | 1.E-02 | -0.4 | 3.E-02 | 0.0 | 1.E+00 | -0.4 | 3.E-02 |
| tpxl-1 | n/a | -2.0 | 1.E-14 | -0.5 | 7.E-02 | -0.2 | 7.E-01 | -0.4 | 2.E-01 |

**Supplemental Figure 8. Bioactivated Tioxazafen Down-Regulates Multiple Categories of Genes Related to the Cell Cycle. A-D.** The log fold-change (LogFC) and FDR for the indicated categories.

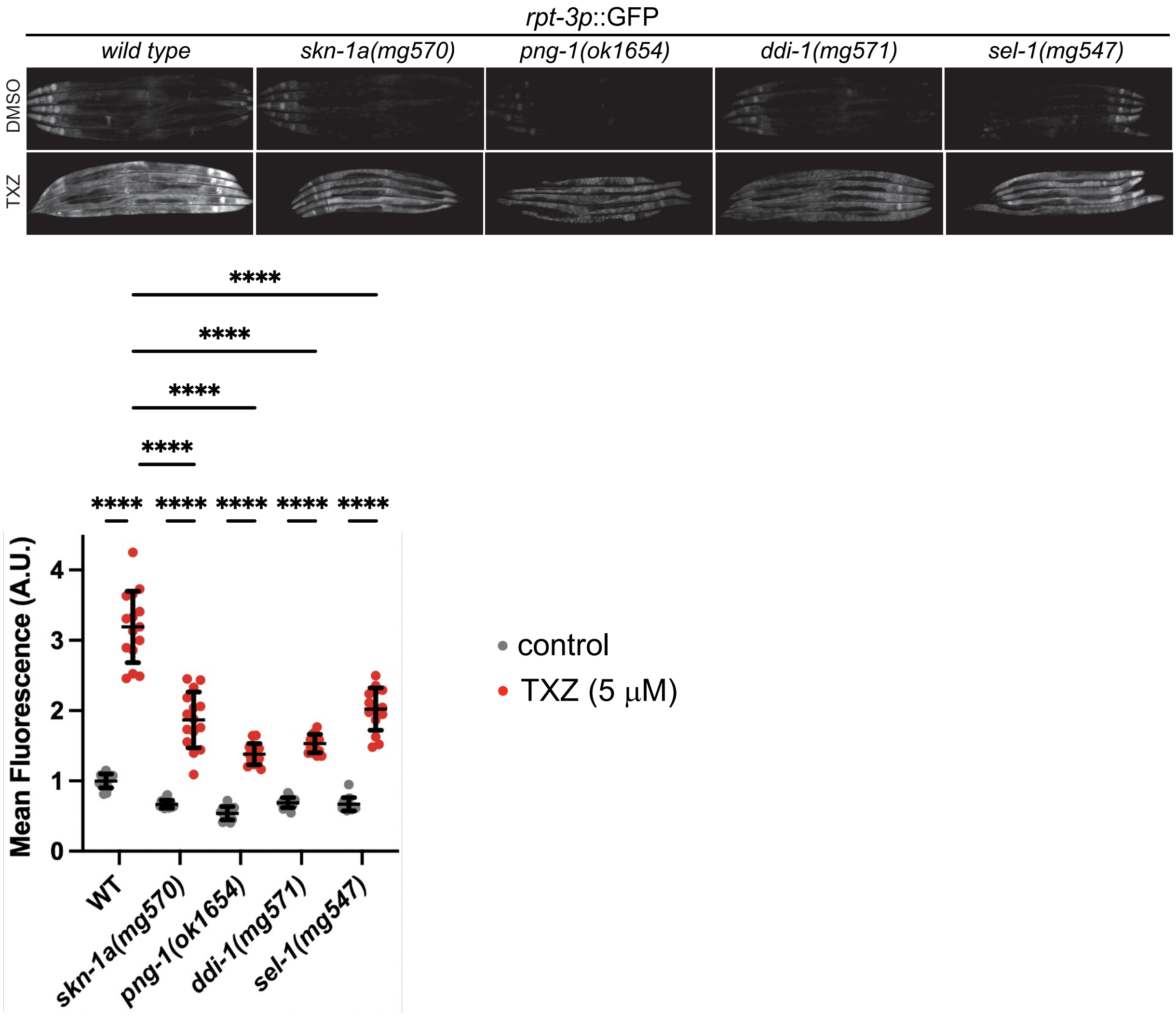


**Supplemental Figure 9. Tioxazafen-Induced Upregulation of the Proteasome Is Only Partially-Dependant on the SKN-1A Pathway.** L4 animals were shifted to plates supplemented with 5 μM tioxazafen or DMSO control and incubated at 20°C for 16 hours before imaging. Quantification of n=15 animals per condition is shown. Error bars show mean ±SD. Significance was determined by ordinary two-way ANOVA with Šidák's multiple comparisons test (**** $p<0.0001$).

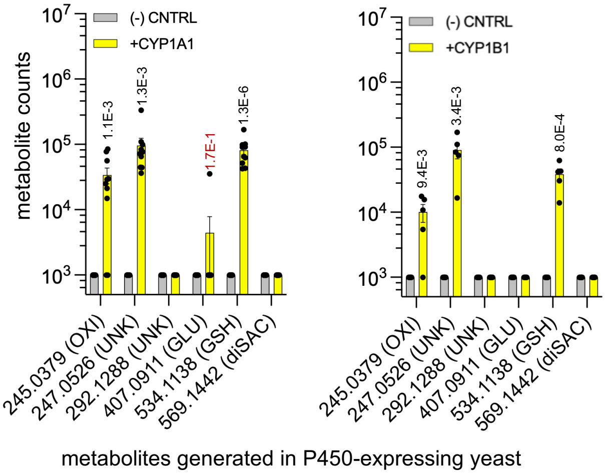


**Supplemental Figure 10. Tioxazafen is Bioactivated by human cytochrome CYP1A1 and CYP1B1.** Tioxazafen-derived metabolites generated in yeast expressing the indicated P450. N≥4, n=2. The Students' T-test *p* values shown atop each bar is in comparison to the (-) control. *p* values are coloured red if $p<0.05$. Exact masses are shown on the x-axis along with the likely type of conjugate in brackets (OXI, oxidized; UNK, unknown; GLU, glucosidated; GSH, glutathione; diSAC, disaccharide). In all graphs, the standard error of the mean is shown.

### Supplemental Figure 11

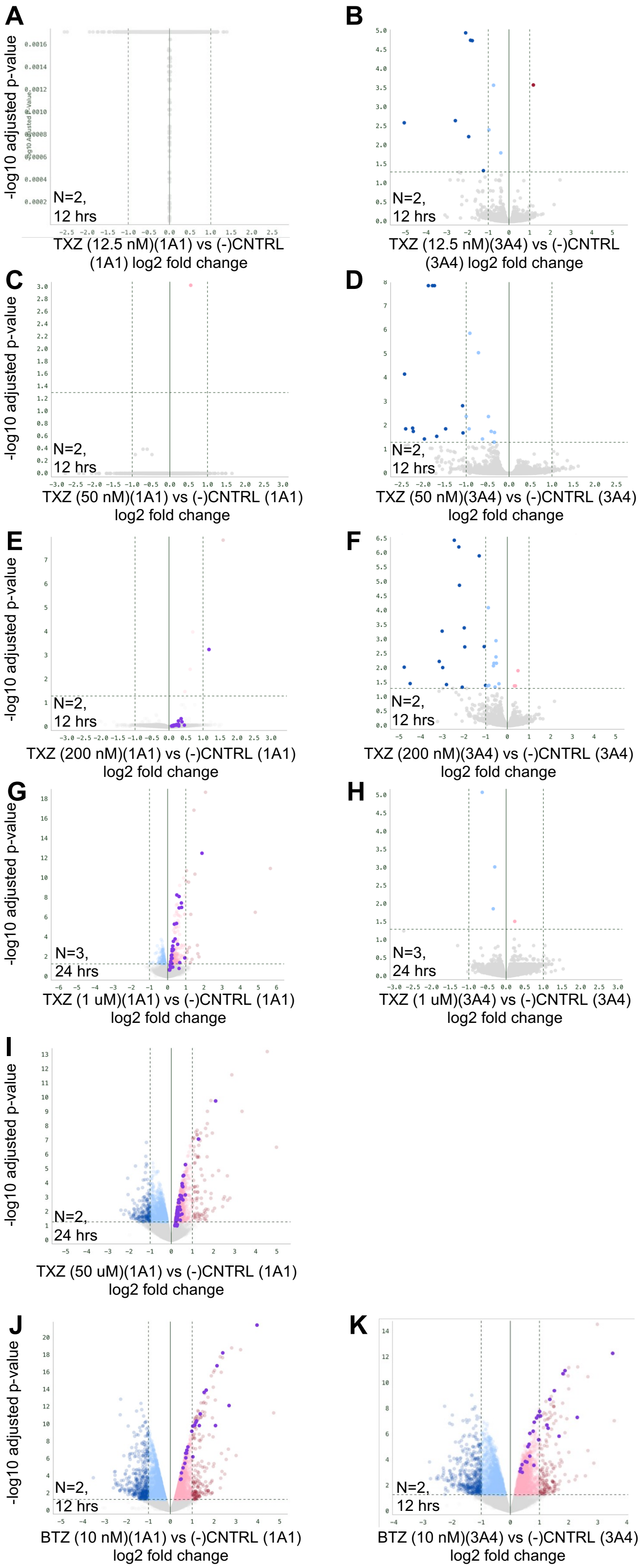


**Supplemental Figure 11. Tioxazafen is Bioactivated by human cytochrome CYP1A1 and CYP1B1. Plot of b**ulk RNAseq analyses showing adjusted p-values as a function of fold change. Shown is data from CYP1A1-expressing HEK293T cells (left) or CYP3A4-expressing control HEK293T cells (right) with the indicated compound at the indicated compound concentration. The number of biological repeats and the incubation time is shown on the inset. Datapoints highlighted in purple are Human Hallmark 2025 genes that belong to the unfolded protein response group.

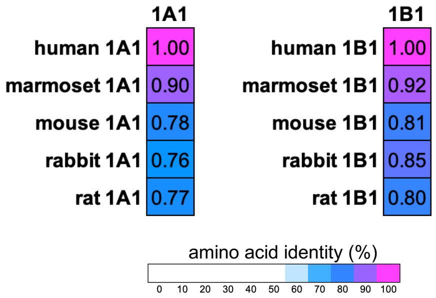


**Supplemental Figure 12.** A comparison of the % sequence identity of the indicated P450s.
